# TWAS pathway method greatly enhances the number of leads for uncovering the molecular underpinnings of psychiatric disorders

**DOI:** 10.1101/373050

**Authors:** Chris Chatzinakos, Donghyung Lee, Na Cai, Vladimir I. Vladimirov, Anna Docherty, Bradley T. Webb, Brien P. Riley, Jonathan Flint, Kenneth S. Kendler, Nikolaos Daskalakis, Silviu-Alin Bacanu

## Abstract

Genetic signal detection in genome-wide association studies (GWAS) is enhanced by pooling small signals from multiple Single Nucleotide Polymorphism (SNP), e.g. across genes and pathways. Because genes are believed to influence traits via gene expression, it is of interest to combine information from expression Quantitative Trait Loci (eQTLs) in a gene or genes in the same pathway. Such methods, widely referred as transcriptomic wide association analysis (TWAS), already exist for gene analysis. Due to the possibility of eliminating most of the confounding effect of linkage disequilibrium (LD) from TWAS gene statistics, pathway TWAS methods would be very useful in uncovering the true molecular bases of psychiatric disorders. However, such methods are not yet available for arbitrarily large pathways/gene sets. This is possibly due to it quadratic (in the number of SNPs) computational burden for computing LD across large regions. To overcome this obstacle, we propose JEPEGMIX2-P, a novel TWAS pathway method that i) has a linear computational burden, ii) uses a large and diverse reference panel (33K subjects), iii) is competitive (adjusts for background enrichment in gene TWAS statistics) and iv) is applicable as-is to ethnically mixed cohorts. To underline its potential for increasing the power to uncover genetic signals over the state-of-the-art and commonly used non-transcriptomics methods, e.g. MAGMA, we applied JEPEGMIX2-P to summary statistics of most large meta-analyses from Psychiatric Genetics Consortium (PGC). While our work is just the very first step toward clinical translation of psychiatric disorders, PGC anorexia results suggest a possible avenue for treatment.

## 1 Introduction

Genome-wide association studies (GWAS) have been very successful for identifying diseases loci using single-marker based association tests (Bush & Moore, 2012). Unfortunately, such methods have had limited power to identify causal genes or pathways (Wang, Li, & Hakonarson, 2010). For most complex traits, genetic risks are likely the result of the joint effect of multiple genes located in causal pathways (Ramanan, Shen, Moore, & Saykin, 2012). Consequently, pooling information across genes in a pathway is likely to greatly improve signal detection.

Given that gene expression (GE), is widely posited to be the critical causal mechanism linking variant to phenotype (Emilsson et al., 2008), the pooling of information across multiple variants should be mediated by GE. Gene expression based methods, widely denoted as transcriptome wide association analysis (TWAS), exist for gene-level inference (Chatzinakos et al., 2017; Gamazon et al., 2015; Gusev et al., 2016). They combine summary statistics at expression Quantitative Traits Loci (eQTL) known to best predict GE to infer the association between trait and GE for the gene under investigation. While TWAS pathway methods do not exist, such methods would help tremendously the translation of psychiatric disorders by i) eliminating the confounding effect of linkage disequilibrium (LD) on TWAS gene statistics and ii) increasing the power by directly modelling the LD between TWAS summary statistics. However, such methods are not yet available, likely due to the large storage/computational burden associated with computing the LD between the numerous SNPs, possibly, involved in computing TWAS statics for numerous genes in a large region, e.g. Major Histocompatibility (MHC) region from chromosome 6p (∼25-35Mbp) or even entire chromosome arms. The large computational burden is the results of the variance of the linear combinations for TWAS statistics (Jin et al., 2014) being assessed using the estimated pairwise LD for all eQTL SNPs which, for *m* variants, requires a heavy *0*(*m*^2^) computational burden.

Currently, pathway analysis methods are non-transcriptomic, i.e. at a minimum they do not use the LD between TWAS gene statistics. Most of them just search for “agnostic” (i.e. not GE mediated) signal enrichment in a pathway/gene set. Among existing pathway methods we mention ALIGATOR (Holmans et al., 2009), GSEA (Subramanian et al., 2005), DAPPPLE (Rossin et al., 2011), as MAGENTA (Segre et al., 2010), INRICH (P. H. Lee, O’Dushlaine, Thomas, & Purcell, 2012) and MAGMA (de Leeuw, Mooij, Heskes, & Posthuma, 2015), as well as online tools: GeneGo/MetaCore (www.genego.com), Ingenuity Pathway Analysis (www.ingenuity.com), PANTHER (www.pantherdb.org), WebGestalt (bioinfo.vanderbilt.edu/webgestalt), DAVID (david.abcc.ncifcrf.gov) and Pathway Painter (pathway.painter.gsa-online.de. While not designed for pathway analyses, LDpred (Bulik-Sullivan et al., 2015; Finucane et al., 2015) can also be adapted to test whether pathways are enriched above the polygenic background while adjusting for genomic covariates. Although all these tools were shown to be very powerful, TWAS based pathway analyses can greatly complement the “agnostic” findings of all these tools.

To advance the translation of psychiatric disorders we propose JEPEGMIX2 Pathway (JEPEGMIX2-P) that extends the reach of TWAS methods to pathway-level. It promises to increase power by directly modeling the LD between TWAS gene statistics. JEPEGMIX2-P i) uses a very large and diverse reference panel consisting of 33K subjects (including around >10K Han Chinese), ii) automatically estimates ethnic composition of cohort, iii) uses these weight to compute LD for gene statistics via a linear running time procedure, iv) uses LD and GWAS summary statistics to rapidly test for the association between trait and expression of genes even in the largest pathways, v) is competitive, i.e. adjusts for the background enrichment of TWAS gene statistics, and vi), to avoid the large signal in a gene inducing significant signals in all small pathways that include it, provides the option of a conditional analysis that eliminates the effect of SNPs with significant signals. When compared to MAGMA, its analyses of PGC GWAS data yield a markedly increased number of significant signals.

## 2 Methods

Naïve application of many analysis methods comparing the statistic with the default null hypothesis (*H*_0_), when applied for genes/pathways with numerous SNPs/genes might yield large signals merely by accumulating “average” polygenic signals from well-powered studies. This comparison to the default null is also known as uncompetitive statistic, as it does not take into account the average enrichment of the genome. To avoid such an accumulation of average polygenic information in the uncompetitive statistic, we use competitive tests that adjust the SNP and gene level *Χ*^2^ statistics for the background enrichment of genome wide SNPs and TWAS gene statistics, respectively. This is achieved simply by adjusting gene statistics for average non-centrality (Text S1, S2 of Supplementary Information-SI). Subsequently, as detailed in S3 and S4] we use the GWAS summary statistics i) to estimate the ethnic composition of the study cohort and ii) use the estimated ethnic weights to build a pathway statistic that has a highly desirable *O*(*m*) computational burden.

GWAS summary data comprise of a large range of effect sizes and it is unclear whether the estimated pathway statistics are related to the whole range, including SNPs with very small effects, or just SNPs with very large effects. To avoid a very large signal in a gene inducing a significant signal in all smaller pathways including the gene, we also offer the option to eliminate the effect of SNPs with statistically significant signals, by applying a novel conditional analysis procedure (Text S5 in SI) to summary statistics before their use in our TWAS pathway tool.

### 2.3 Computation of pathway statistic

Generic TWAS methods, including our JEPEGMIX2m, output Z-score statistics by gene. Thus, if the correlation between gene statistics is available, e.g. by using the *0*(*m*) method described above, these statistics can be combined using a Mahalanobis *Χ*^2^ statistics with the number of degrees of freedom (df) equal to the number of genes. Unfortunately, this can become quickly very involved if we need to compute the LD between statistics of all ∼20,000 genes. However, given that the genotypes of variants in different chromosome arms are practically independent, if follows that Z-scores for genes on different chromosome arms are independent. Thus, the Mahalanobis type statistic can be computed more easily by i) computing chromosome arm *Χ*^2^ statistics and, subsequently, ii) combining the resulting chromosome arm statistic in a *Χ*^2^ pathway statistic (Fig 1). Similarly, the df for the *Χ*^2^pathway statistic equals the sum of the dfs for chromosome arm statistics.

**Fig 1.**
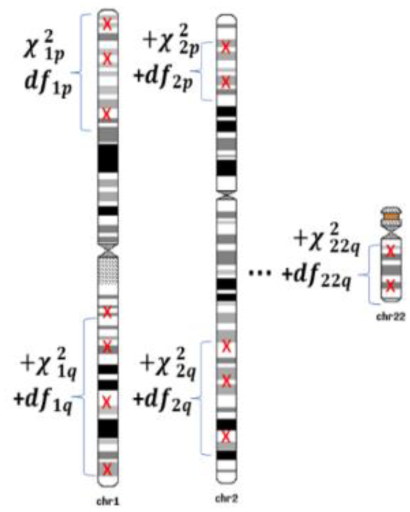
Computation of pathway statistics.

### 2.4 Annotation and reference panel

As pathway database we used MSigDB (Liberzon, 2014; Liberzon et al., 2015; Liberzon et al., 2011), which is well maintained and widely used by researchers. To facilitate user-specific input for new pathways along with future extensions, the annotation file for JEPEGMIX2-P now includes an R-like formula for the expression of each gene as a function of its eQTL genotypes and of the content for each pathway as a function of the names its constituting genes. The updated annotation file includes cis-eQTL for all tissues available in the v0.7 version of PredictDB (http://predictdb.hakyimlab.org/). To avoid making inference about genes poorly predicted by SNPs, for the 48 available tissues (Text S6, Table S1 of SI), we retain only genes for which the expression is predicted with reasonable accuracy [False Discovery Rate (FDR) q-value < 0.05] from its eQTLs. The current version uses the 32,953 subjects (33K) as the reference panel. It consists of 20,281 Europeans, 10,800 East Asians (from CONVERGE study, Text S5 of SI), 522 South Asians, 817 Africans and 533 Native of Americas (Text S6 Table S2 of SI).

## 3 Simulations

To estimate the false positive rates of JEPEGMIX2-P, for five different cosmopolitan studies scenarios, we simulated (under *H*_0_) 100 cosmopolitan cohorts of 10,000 subjects for Ilumina 1M autosomal SNPs using 1KG haplotype patterns (D. Lee et al., 2015) (Text S9, Table 3 of SI). The subject phenotypes were simulated independent of genotypes as a random Gaussian sample. SNP phenotype-genotype association summary statistics were computed from a correlation test. For each cohort, we obtained JEPEGMIX2-P statistics, for the two “null” enrichment scenarios i) under null (*H*_0_), of no enrichment, and ii) polygenic null (*H*_*p*_), i.e. when enrichment is uniform over the entire genome regardless of functionality of individual genomic regions. For the JEPEGMIX2-P analyses of the resulting data we used i) prespecified (PRE) and ii) automatically estimated ethnic weights (EST). The prespecified (PRE) weights were assigned assuming information from the studies about subpopulations involved were available. As PRE-weights, we assigned “study-published” weights for the closest subpopulations from our new reference panel. Given that i) subjects were re-assigned to subpopulations in the new panel and ii) the populations labels in the new panel do not correspond to the ones from 100 Genomes, this induced possible mismatches that might result in increased false positive rates. To avoid this, a second version of the PRE-approach provides the published weights to continental superpopulations, i.e. EUR, ASN, SAS, AFR and AMR.

During our initial simulations we observed that pathways with name lengths ≤ 8, e.g. pathways denoting chromosome bands like: chr3p21, ch6p21 etc., have increased false positive rates due to having numerous genes in high LD because of their proximity. For that reason, we also estimated the size of the test for all cohort scenarios just for these high LD pathways.

Recently, due to its strength, MAGMA is one of the most used pathway analysis methods. Consequently, we compare the results obtained from our method with those obtained by this state-of-the-art method. However, to compare JEPEGMIX2-P with MAGMA, given that 1) the simulated cohorts might not reflect real data and 2) these data sets are for cosmopolitan cohorts (MAGMA software does not provide reference panel for these cohorts), we used real data to create “nullified” data sets. These nullified data sets are based on 20-real GWAS (of mostly Caucasian cohorts) like: schizophrenia (SCZ), attention deficit hyperactive disorder (ADHD), autism (AUT), major depressive disorder MDD (see Table 1) and a further (preponderantly European) 16 data sets that are not yet publicly available. This approximation for null data is obtained by substituting the expected quantile of the Gaussian distribution for the (ordered) Z-score (see also Text S1, nonparametric robust estimation of weights section, of the SI) after eliminating SNPs with significant association p-values in the original GWAS. However, one side effect of this approach consists of statistics within/near the peak signals in original GWASs might be still slightly more concentrated into the tails of the distribution to be a perfect “null data". This can result in a slight increase in false positive rate, especially when applied to the nullified version of a highly enriched GWAS (e.g. PGC2 SCZ). However, most of the data sets used in “nullification” were not highly enriched in association signals.

**Table 1.** Description of GWAS studies and traits that were analyses.

| Trait | Trait Abbreviation | Dataset Description |
| --- | --- | --- |
| Schizophrenia | SCZ | PGC2 SCZ (Ripke et al., 2013) |
| Attention Deficit Hyperactivity Disorder | ADHD | PGC ADHD (Demontis et al., 2017) |
| Autism | AUT | PGC AUT (Buxbaum et al., 2012) |
| Bipolar | BIP | PGC BIP (Stahl et al., 2019) |
| Eating Disorders (Anorexia) | ED | PGC EAT (Duncan et al., 2017) |
| Major depression disorder | MDD | PGC MDD (Wray et al., 2018) |
| Post-traumatic stress disorder | PTSD | PGC PTSD (Nievergelt et al., 2019) |

### 3.1 Practical Applications

We applied JEPEGMIX2-P and MAGMA to summary statistics coming from Psychiatric Genetics Consortium (PGC-http://www.med.unc.edu/pgc/) datasets, i.e. Schizophrenia (SCZ), Attention Deficit Hyperactivity Disorder (ADHD), Autism (AUT), Bipolar (BIP), Eating Disorders (Anorexia) (ED), Major depression disorder (MDD) and Post traumatic stress disorder (PTSD) (see Table 1). To limit the increase in Type I error rates of JEPEGMIX2-P, we deem as significantly associated only those pathways that yield an FDR-adjusted p-value (q-value)< 0.05. Due to *C4* explaining most of Major Histocompatibility (MHC) (chr6:25-33 Mb (McCarthy et al., 2016), gene/signals for SCZ, for this trait, we omit non-*C4* genes in this region. Moreover, due to the high correlation between SNPs in MHC (chr6:25-33 Mb), we also omit genes in this region for MDD, which also showed MHC signals (Wray et al., 2018).

## 4 Results

JEPEGMIX2-P using our proposed automatic weight detection procedure (see Methods), controlled the false positive rates at or below the nominal threshold, even when this threshold was 10^−6^, under both null (*H*_0_) and “polygenic null” scenario (*H*_*p*_ - enrichment in association signals is uniform over the entire genome, in keeping with the polygenicity of most traits). When the method used narrowly prespecified subpopulation weights (e.g. using the closest subpopulations from the reference sample, i.e. as derived from the study description), the false positives rates were increased, especially for lower nominal rates, by up to ∼220-450 (Text S9, Figs S1-S5 in SI). However, JEPEGMIX2-P with pre-estimated weights based on super populations (i.e. European, East Asian, African etc.) had a much lower inflation of false positive rates; only for 10^−6^ threshold the false positive rate was increased by ∼2-4 times, under both *H*_0_ and *H*_*p*_ scenarios (Text S9, Fig S6 in SI).

For high-LD pathways, e.g. those defined by single chromosome bands in MSigDB (Liberzon, 2014; Liberzon et al., 2015; Liberzon et al., 2011), the behavior of JEPEGMIX2-P with automatically estimated weights is similar to the one for the whole set of MSigDB pathways. However, false positive rates increase by ∼300-1,200 for the narrowly prespecified subpopulation weights (Text S9, Figs S7-S11 in SI), while when using super population-based weights, it remained practically unchanged from the 2-4X increase derived for all pathways (Text S9, Fig S12 in SI).

Using the FDR p-value adjustment, for both unconditional and the very conservative conditional JEPEG2-P analyses, we uncovered numerous significant pathway signals for Psychiatric Genetics Consortium (PGC) traits (Table 1). For the most significant we present heatmaps (Fig. 2-5, Text S10, Fig. S13-S19 in SI) while extended tables (Supplementary Excel file) include all significant signals. On the other hand, most likely due to not modelling the LD between gene statistics, MAGMA found fewer signals when it was applied to the TWAS gene statistics from JEPEGMIX2 (Table 2). (The number of MAGMA signals was even lower when inputting SNP GWAS instead of gene TWAS summary statistics.) JEPEGMIX2-P running time for a TWAS gene and pathway analysis of PGC data was less than 5 days on a single core of a cluster node with 4x Intel Xeon 6 core 2.67 GHz.

**Table 2.** Numbers of signals found by JEPEGMIX2-P and MAGMA.

| Trait | JEPEGMIX2-P without conditional analysis | JEPEGMIX2-P conditional analysis | MAGMA |
| --- | --- | --- | --- |
| ADHD | 1 | 1 | - |
| AUT | 2 | 2 | - |
| ED | 268 | - | - |
| MDD | 607 | 7 | - |
| SCZ | 825 | 27 | 42 |
| PTSD | - | - | - |

**Fig 2.**
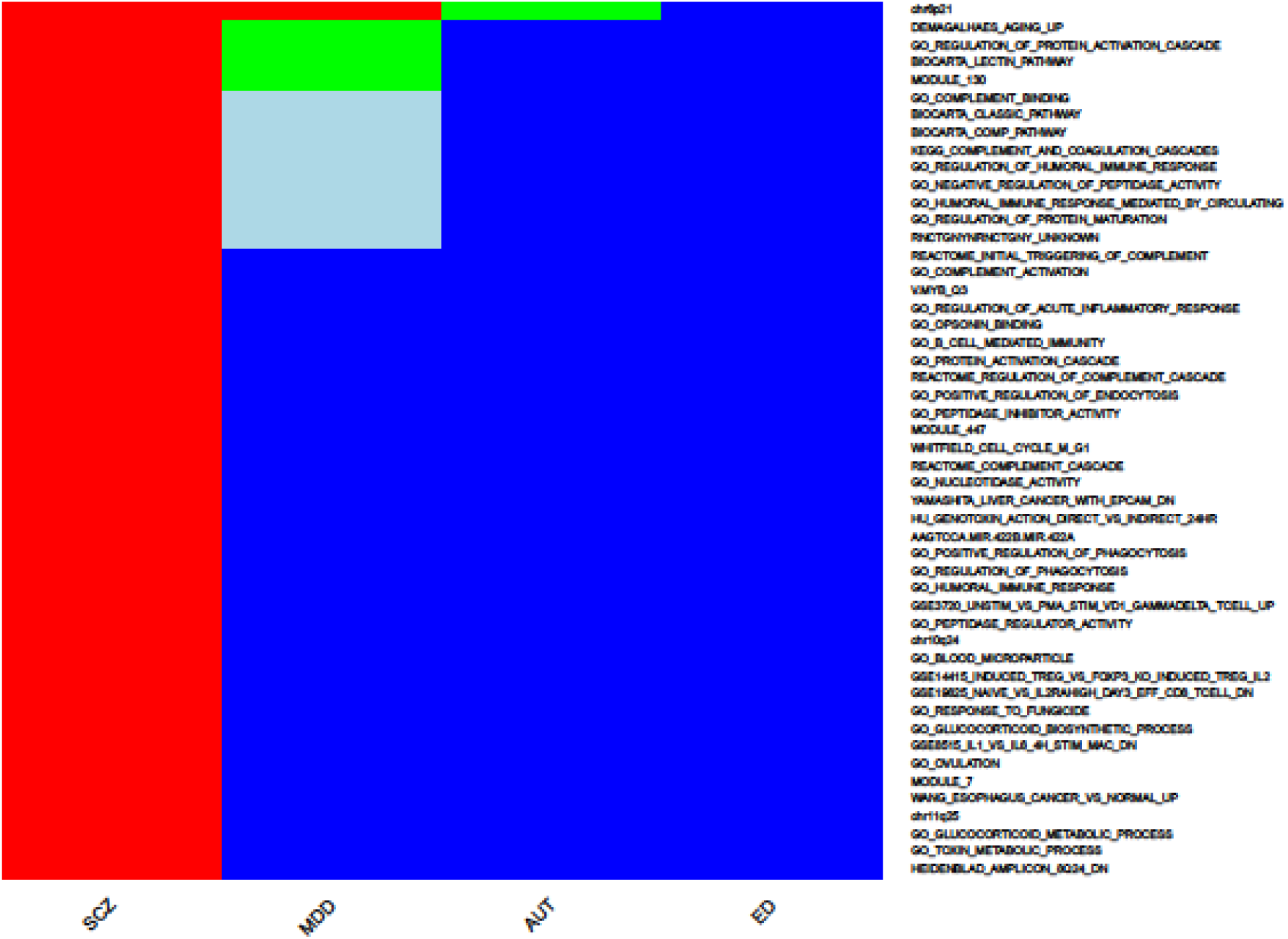
Top 50 pathways for psychiatric traits (without conditioning out significant SNP signals). Where red color denotes *q* < 0. 001, orange 0. 001 < *q* < 0. 01, green 0. 01 < *q* < 0. 05, light blue 0. 05 < *q* < 0. 16 and blue 0. 16 < *q* < 1.

**Fig 3.**
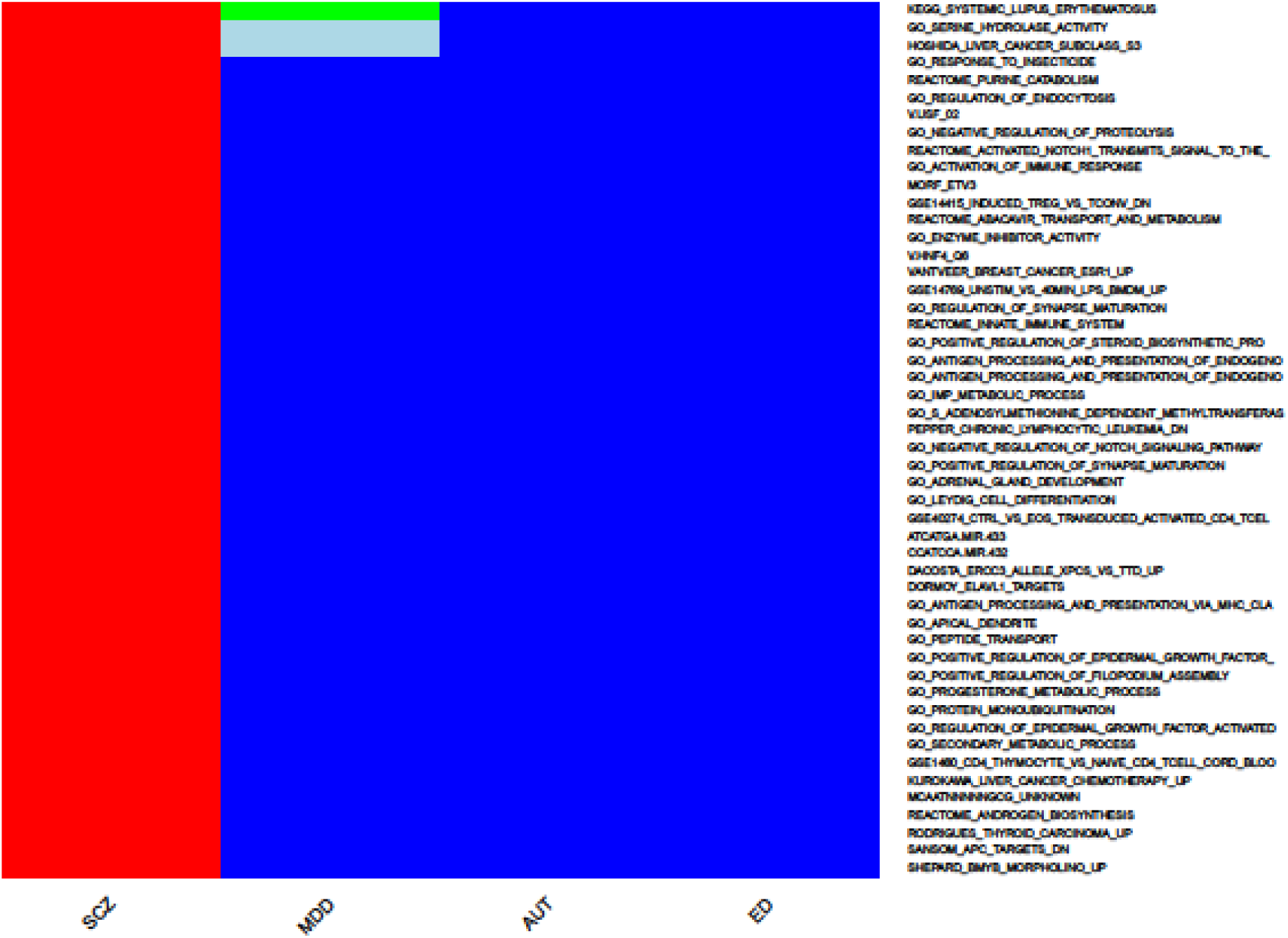
Top 51-100 for psychiatric traits (without conditioning out significant SNP signals). See Fig S2 for background.

**Fig 4.**
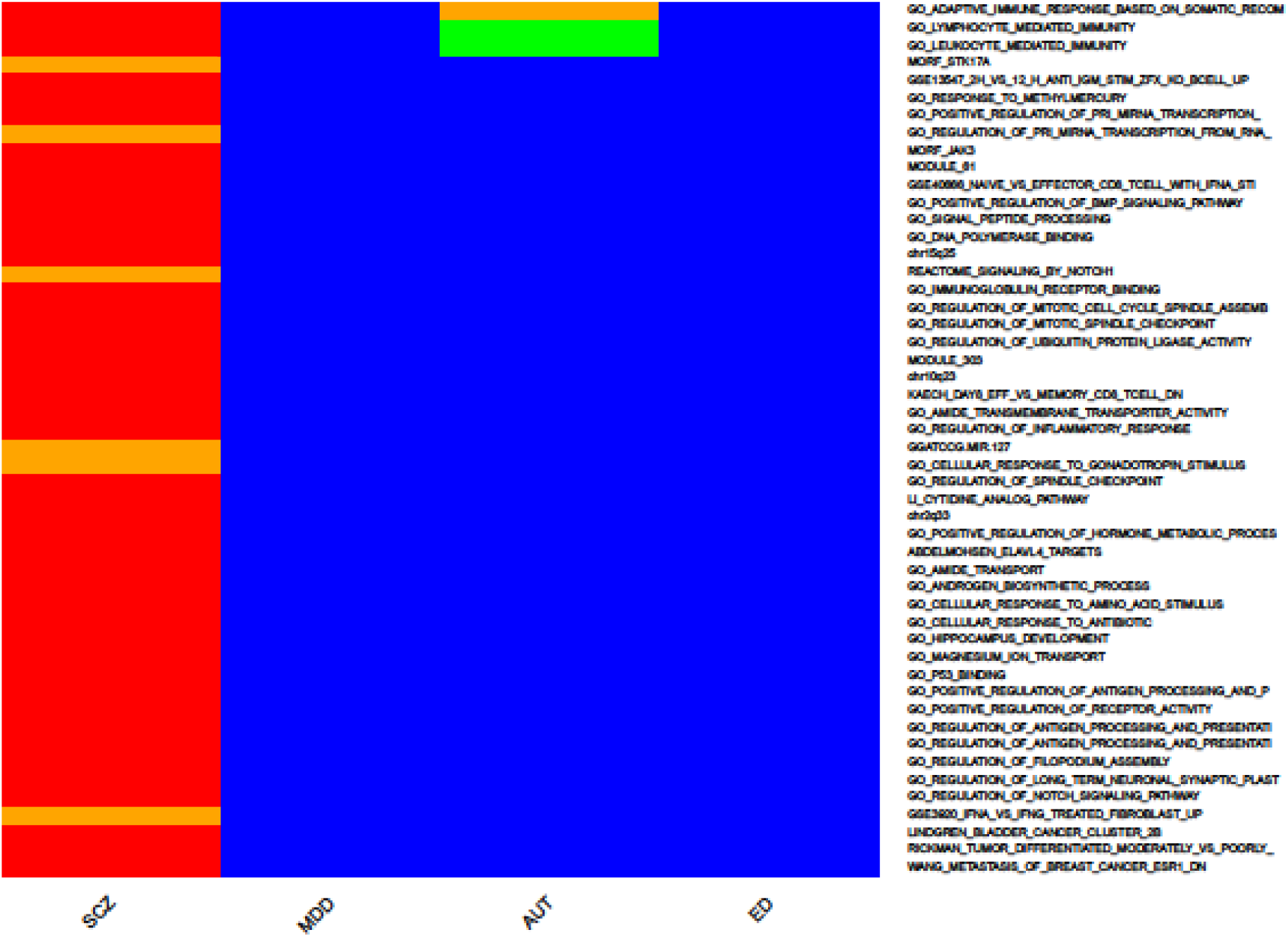
Top 101-150 for psychiatric traits (without conditioning out significant SNP signals). See Fig S2 for background.

**Fig 5.**
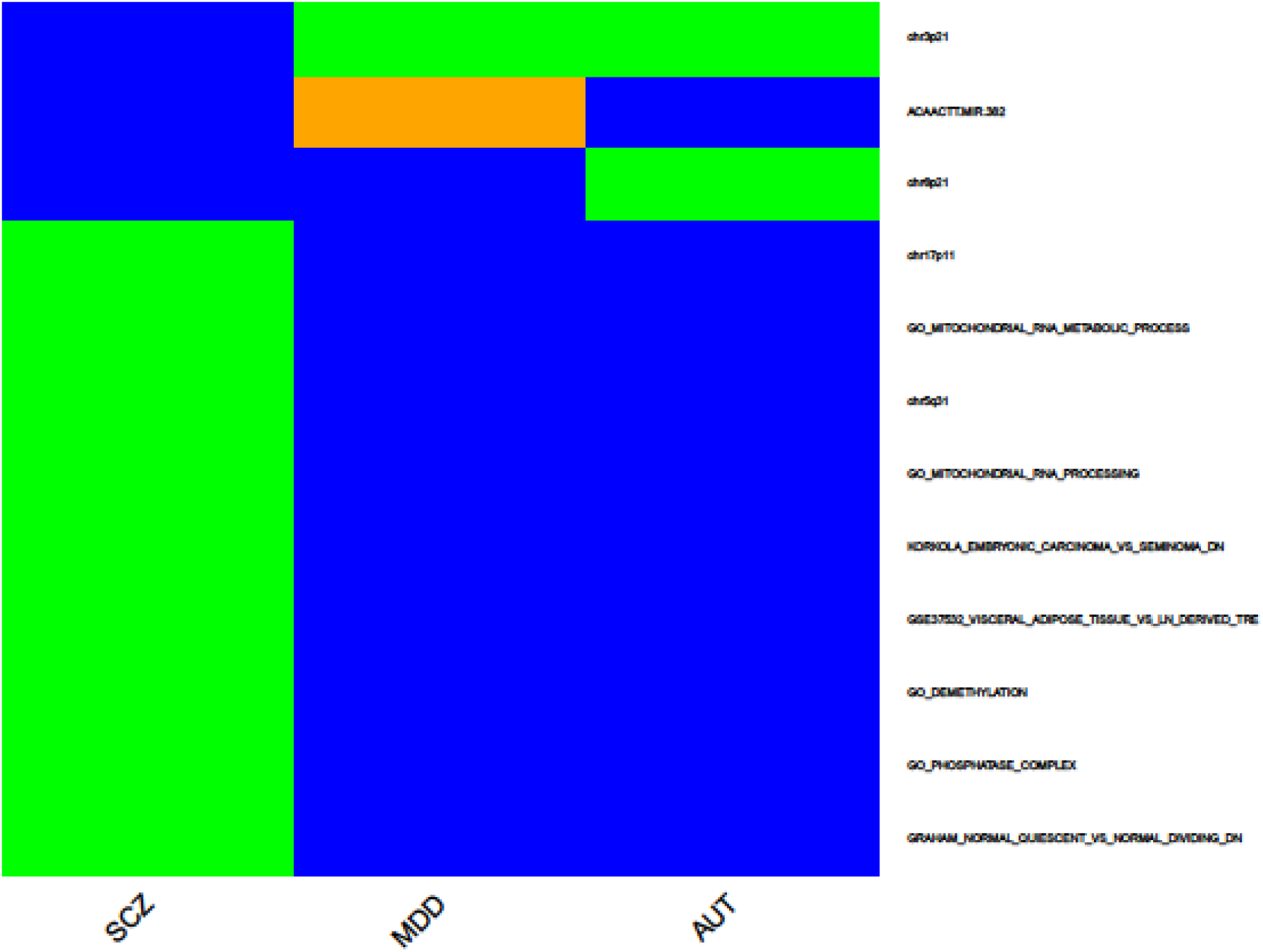
Top pathways for psychiatric traits when conditioning out significant signals. See Fig S2 for background.

## 5 Discussion

The discovery of biological pathways implicated in diseases is the target for any genetic analysis. Despite the numerous methods available for pathway analyses, none of these methods relies solely on eQTLs to infer the association between expression of genes in pathway and trait, which is widely posited to be the critical causal mechanism. To overcome these two main factors, we propose JEPEGMIX2-P for testing the association between pathway expression and trait. Even for enriched GWAS and high LD pathways, JEPEGMIX2-P with the automatic weights fully controls the false positive rates at or below nominal levels.

Narrowly assigning the ethnic weights to the subpopulations perceived as the “closest” to the ones in the studies is not advisable due to the possibility of great mismatch between the cohort and the “re-arranged” subpopulations from our reference panel, which can result in greatly increased false positive rates (see Methods). Consequetly, users should use the automatic detection of cohort composition, regardless whether cohort allele frequencies are available or not.

While the method is a welcome addition to our pathway tools, it still has major limitations when attempting to use it for assigning “causal” tissues/cell. First, due to the rather small sample sizes of existing GE experiments ∼80% of genes do not have good GE prediction from eQTL SNPs. Second, there is a large difference between the sample sizes available for different tissues; generally, tissues that are more accessible have larger sample sizes.

However, we want to stress that, both at present and in the future, from all practical TWAS analyses the most important findings are the pathways and genes that are associated with the trait, not the tissue/single cell where they are (the most) significant. This argument is strongly supported by a very recent review in Nature Reviews Genetics (Hekselman & Yeger-Lotem, 2020); it shows that while mis-regulation of gene is causal in one/few tissues, these genes are mis-regulated in numerous more tissues. Consequently, even if we do not assay the gene/pathway in the causal cell types directly, we can find the mis-regulated genes by detecting signals in many other tissues and cell types.

Applying JEPEGMIX2-P to psychiatric phenotypes, we discovered numerous pathways that were deemed significant for SCZ, ADHD, AUT, EAT and MDD. We mention that while the original SCZ paper did not report any genome-wide significant pathways and MAGMA reports only five (and, in our analysis 42, after inputting TWAS gene statistics), JEPEGMIX2 detected hundreds of them. Even more, these signals are not very likely to be false discoveries due to our method i) being competitive, i.e. adjusting both SNP and gene statistics for polygenic/gene enrichment background, ii) accurately estimating of ethnic weights and iii) excellent control of Type I error rates.

Interpreting and validating all pathway signals require substantially more work. It is always “the last mile” that is the most laborious part of the translation process. However, JEPEGMIX2-P provides carefully vetted targets for wet-lab validation (see Supplementary Excel Spreadsheet). Nonetheless, our findings sometimes allow for some reasonably informed inferences. For instance, in anorexia results, the most significant signal in a tissue correspond to GEISS_RESPONSE_TO_DSRNA_UP pathway (Supplementary Excel file), which is a pathway that is involved in response to virus infections. This finding suggests a possible avenue of treatment that is easily available and inexpensive to try: anorexia patients with active virus infections might benefit from being treated with suitable anti-viral medication. However, the responders to such treatment are likely to form only a minority of the anorexia patients, i.e. a fraction of those with active viral infections that are treatable by drugs.

