## Supplemental Material for "TWAS pathway method greatly enhances the number of leads for uncovering the molecular underpinnings of psychiatric disorders"

### S1. Centralize SNP Z-scores for computing a competitive gene level statistic

To avoid an accumulation of just averagely enriched polygenic variant information, we competitively adjust SNP $\chi^2$ statistics for background enrichment. This is achieved by adjusting the statistic for average non-centrality.

Let $Z$ be the vector of $Z$-scores for measured SNPs in the genome scans. Due to polygenicity, the expected genome scan $\chi_1^2 = Z^2$ statistics, each with 1 degree of freedom (df), has a non-zero background noncentrality parameter $\lambda^2$, i.e. $E(Z^2) = 1 + \lambda^2$. Thus, by the method of moments, we can estimate $\widehat{\lambda^2} = \overline{Z^2} - 1$, where $\overline{Z^2}$ is computed using all measured SNPs in the genome scan, However, given that $\lambda^2 \geq 0$ , a better estimator is, thus, $\widehat{\lambda^2} = \max(\overline{Z^2} - 1, 0)$. To develop a competitive test, before computing gene-level statistics, Z-scores must be shrunk towards zero by adjusting for the average background enrichment ($\widehat{\lambda^2}$). This can be achieved via a 3-step process:

(1) Recompute, under "average" noncentrality, the p-value associated with $\chi_1^2$ statistics: $P' = 1 - F(Z^2 | \widehat{\lambda^2})$, where $F(.| \widehat{\lambda^2})$, is the cumulative distribution function (cdf) of the non-central $\chi_1^2$ distribution with 1 df and noncentrality parameter $\widehat{\lambda^2}$.

(2) Transform this p-values vector, $P'$, into its quantile vector from a central $\chi_1^2$ distribution with 1 df, i.e. $\chi^2 = F^{-1}(1 - P' | \lambda^2 = 0)$,

(3) Transform $\chi^2$ to a "central" Z-score: $Z' = sign(Z) * \sqrt{\chi^2}$.

By Delta method (a first-order Taylor approximation), $Z'$ is a linear transformation (deflation) of $Z$ and it has the same correlation structure. Thus, $Z'$ can be used to build the competitive statistics, which exhibit a variance that is identical to their associated non-competitive versions.

### Nonparametric robust estimation of weights

To estimate robust weights and to avoid false positives we apply a two-step, robust algorithm to the $Z$-scores of the SNPs. **First,** let $Z_\sigma = (z_{\sigma_1}, z_{\sigma_2}, \dots, z_{\sigma_m})$, where $\sigma$ indicates the permutation of indices of Z-scores, $Z$, for the $m$ SNPs, that orders these statistics in increasing order. **Second 2)** $z_i' = \Phi^{-1}(\frac{\sigma_\iota}{m+1})$, where $\Phi^{-1}$ is the inverse normal cumulative distribution function. Subsequently, these transformed risk scores are used for computing ethnic weights.

### S2. Centralize gene level Z-scores for computing pathway statistics

Coding regions (e.g. genes), due to their functionality, are expected to be enriched above the average polygenic background of the genome. Thus, when computing pathway statistics, to avoid an accumulation of just averagely enriched coding gene information, we competitively adjust transcriptomic gene statistics for background enrichment. This is achieved by adjusting the$\chi^2$/Z-score gene statistics for their non-centrality. Consequently, prior to using in competitive pathway statistics, the gene Z-scores are centralized using a process similar to S1 (Nonparametric robust estimation of weight section) but applied to gene instead of SNP statistics.

### S3. Automatic detection of the ethnic composition for the cohort.

The LD between markers can vary widely between human populations. Thus, to compute the LD, which is necessary for internal imputation and variance estimation for gene statistics, we need to estimate the ethnic composition of the cohort. Our group has previously described, in DISTMIX paper [1], a method of using the reference panel to estimate the ethnic composition when the cohort allele frequencies (AF) are available. However, lately consortia do not provide such summary measure; they often might provide just the Caucasians AF. *Consequently, there is a need for a method to estimate the ethnic composition of the cohort even when no AFs are provided*. Below is the theoretical outline of such method, which uses only the SNP summary statistics (Z-scores).

Assume that the cohort genotype is a mixture of genotypes from $k$ ethnic subpopulation from a large and diverse reference panel. If the $i$-th subject at the $j$-th SNP has genotype $G_{ij}$ and belongs to the $l$-th group, let $p_j^{(l)}$ be the frequency of the reference allele frequency for this SNP in the $l$-th group. Let $q_j^{(l)} = 1 - p_j^{(l)}$ and $G'_{ij} = \frac{G_{ij} - 2\,p_j^{(l)}}{\sqrt{2p_j^{(l)}\,q_j^{(l)}}}$ be the normalized genotype, i.e. the transformation to a variable with zero mean and unit variance. Near $H_0$, as outlined in Lee at al. [2], SNP Z-score statics $Z_j s$ have the approximately the same correlation structure as the genotypes used to construct it, $G_{*j}$'s. Given that $G'_{*j}$ is a simple linear transformation of $G_{*j}$ with a positive slope, by the basic properties of the correlation function, it follows that $G_{*j}$ and $G'_{*j}$ have the same correlation structure. Consequently, the Z-statistics ($Z_j s$) have the same correlation structure as $G'_{*j}$'s. However, given that both $G'_{*j}{}'s$ and $Z_j s$ have unit variance, it follows that the two have the same *covariance (i.e. not only the same correlation)* structure. It follows that any functional transformation of $Z_j s$ is identically distributed as the same functional transformation of $G'_{*j}{}'s$. Therefore, for any $s \geq 1$ we can write: $E(Z_j\ Z_{j+s}) = E(G'_{*j}\ G'_{*\,j+s})$, which, assuming that $w^{(l)}$ is the expected fraction of subjects from the entire cohort that belong to the $l$-th subpopulation from the reference panel, becomes

$$E(Z_j\ Z_{j+s}) = \sum_{l=1}^{k} w^{(l)}\ E\left[{G'_{*j}}^{(l)}\ {G'_{*(j+s)}}^{(l)}\right] = \sum_{l=1}^{k} w^{(l)}\ Cov\left({G'_{*j}}^{(l)}, {G'_{*(j+s)}}^{(l)}\right) = \sum_{l=1}^{k} w^{(l)}\ Cor({G'_{*j}}^{(l)}, {G'_{*(j+s)}}^{(l)}) \quad (1).$$

Henceforth, we will simply denote the $\boldsymbol{w}$ vector as weights. While $Cor({G'_{*j}}^{(l)}, {G'_{*(j+s)}}^{(l)})$ is unknown, it can be easily estimated using their reference panel counterparts with appropriate ethnic weights. Thus, the weights, $w^{(l)}$, can be simply estimated by simply regressing the product of Z-scores of reasonably close SNP $Z$-scores, $Z_j\ Z_{j+s}$, on correlations between normalized genotypes at the same SNP pairs for all subpopulations in the reference panel. For all simulations and applications, we use $s = 1$. Because some GWAS might have numerous large signals, e.g. latest height meta-analysis [3], a more accurate estimation of the weights in equation (1) is very likely to be obtained by the process of “nullifying the GWAS Z-scores” i.e., substituting the expected Gaussian quantiles for the ordered $Z_j$ (see S1, nonparametric robust estimation of weights section, in SI). Due to the strong LD among neighboring SNPs, the estimation of the correlation using all SNPs in a genome simultaneously might lead to a poor regression estimate in (1). To avoid this, we sequentially split GWAS SNPs into $1000$ non-overlapping SNP sets, e.g. first set consists of the $1$-st, $1001$-st, $2001$-st, etc. map ordered SNPs in the study. The large distances between SNPs in the same set make them quasi-independent which, thus, improves the accuracy of the estimated correlation. $W = (w^{(l)})$ is subsequently estimated as the average of the weights obtained from the 1000 SNP sets. Finally, we set to zero the negatives weights and normalize the remaining weights to sum to $1$ [4]. While approximate continental (European [EUR], East Asian [ASN], South Asian [SAS], African [AFR] and America native [AMR]) ethnic distribution of subjects can be easily estimated from study info, it is not always clear how these weights should be allocated among continental subpopulations. This further apportioning is likely to be important when the GWAS cohorts contain a large number of admixed populations, e.g. African Americans and American native populations, which in the making of the reference panel, had many subjects re-assigned to related subpopulations. Consequently, when continental proportions are provided by the users, we can use the above described automatic detection to distribute these weights to the most likely subpopulations in the reference panel.

### S4. $\boldsymbol{O(m)}$ LD estimation procedure.

It is very computationally challenging [$O(m^2)$] for $m$ genetic variants] to estimate the large correlation matrices needed to compute TWAS pathway statistics (substantially more so for the upcoming larger reference panels). The same heavy computational burden occurs in fine-mapping when there is a desire to output correlation between statistics of genes and pathways with suggestive/significant signals. Thus, for computational feasibility, we need to find an approach that avoids computing correlation matrices. For the theoretical justification of such an approach we use the mathematical notation from the automatic weight estimation, where $G'_{*j} = \frac{G_{ij} - 2\, p_j^{(l)}}{\sqrt{2\, p_j^{(l)} q_j^{(l)}}}$ is the normalized version of $G_{*j}$, i.e. with means $0$ and variance $1$. As mentioned above, under the null hypothesis, for the same variant $G'_{*j}$s have the same distribution as the Z-scores. The Z-score TWASs statistic per gene or pathway is a linear combination of the Z-scores from expression Quantitative Trait Loci (eQTL) SNPs [5]: $Z = \frac{\sum_{j=1}^{m} b_j\, Z_j}{SD(\sum_{j=1}^{m} b_j\, Z_j\,)}$ , where the $SD(\sum_{j=1}^{m} b_j\, Z_j\,)$ is not known and should be estimated reasonably fast. Thus, in general we are interested in computing the covariance between two very large pathway scores (or the variance of a large one), i.e. linear combinations of Z-scores: $Cov(\sum_{j=1}^{m} a_j\, Z_j, \sum_{j=1}^{m} b_j\, Z_j)$. As stated above, working "by SNP" and computing the correlation is $O(m^2)$ an, thus, highly untenable for very large combinations of SNP statistics. However, it is possible to work by "mimicking" the higher order entity (gene, pathways) statistics by observing that, under the null hypothesis, $\sum_{j=1}^{m} a_j\, Z_j$ and $\sum_{j=1}^{m} b_j\, Z_j$ have, due to normalization of $G'_{*j}$, a distribution that is identical to the distribution of $\sum_{j=1}^{m} a_j\, G'_{*j}$ and $\sum_{j=1}^{m} b_j G'_{*j}$ , respectively. Thus, $Cov(\sum_{j=1}^{m} a_j\, Z_j, \sum_{j=1}^{m} b_j\, Z_j) = Cov(\sum_{j=1}^{m} a_j\, G'_{*j}, \sum_{j=1}^{m} b_j\, G'_{*j})$ , which is easily estimated from a reference sample without computing correlation matrices, by using just a highly desirable linear [$O(m)$] running time procedure. For the correlation between two pathway statistics, then: $Cor(\sum_{j=1}^{m} a_j Z_j, \sum_{j=1}^{m} b_j Z_j) = \frac{Cov(\sum_{j=1}^{m} a_j G'_{*j}, \sum_{j=1}^{m} b_j G'_{*j})}{\sqrt{Var(\sum_{j=1}^{m} a_j G'_{*j})}\sqrt{Var(\sum_{j=1}^{m} b_j G'_{*j})}}$ (2)

Within JEPEGMIX-P, the covariances and correlations of the statistics are transparently computed using subject weights reflecting the fraction in the study cohort for each ethnic group from the reference panel. Thus, computing the correlations reduces to simply applying linear combinations to normalized genotype vectors in reference panels followed by very simple estimations of weighted covariance and variance matrices for the two vectors. We need to underscore again that besides the huge memory savings, the proposed method has linear running time while estimating the correlation matrix has a quadratic (in the number of SNPs) running time.

### S5. Conditional Analysis Approach to eliminate the effect size of significant signal SNPs

A single SNPs with very significant signals ($p \ll 5.10^{-8}$) in GWAS data, may induce large signals for many genes and for small(er) pathways that include these genes. To avoid the undue influence of a single signal, we apply a condition out a large signal via a simple algorithm with five steps as described below:

**1)** For each chromosome arm, make a list that includes SNPs with significant signal ($p < 5.10^{-8}$) and eQTL SNPs.

**2)** Find the SNP with the biggest $|Z|$ in the list and compute the LD patterns of the study cohort are estimated as a weighted mixture of the LD matrices for all ethnic groups in a reference panel (see main text), between this SNP and all the other variants in the chromosome arm.

**3)** Compute the conditional $Z$ values by conditioning out the effect of the largest signal ($Z_{max}$) in the list: ${Z_i}^* = \frac{Z_i - \rho_i * Z_{max}}{\sqrt{1-\rho_i^2}}$, where $Z_i$ is the $z$-score for the i-th SNP and $\rho_i$ is the weighted correlation between the SNP with the largest signal and the i-th SNP.

**4)** Apply steps 2-3 until no SNPs in the list yield $p < 5.10^{-8}$.

### S6. Tissues abbreviation of our new annotation data

**Table S1. Tissue Abbreviation.**

| Tissue | Tissue Abbreviation |
|---|---|
| Adipose Subcutaneous | Adip_Subc |
| Adipose Visceral Omentum | Adip_Visc_Om |
| Adrenal Gland | Adr_Gland |
| Artery Aorta | Art_Aorta |

| | |
|---|---|
| Artery Coronary | Art_Coron |
| Artery Tibial | Art_Tib |
| Brain Amygdala | Br_Amygd |
| Brain Anterior Cingulate Cortex BA24 | Br_Antr_cing_cort_BA24 |
| Brain Caudate Basal Ganglia | Br_Caud_bas_gang |
| Brain Cerebellar Hemisphere | Br_Cereb |
| Brain Cerebellum | Br_Cerr_Hemisph |
| Brain Cortex | Br_Cort |
| Brain Frontal Cortex BA9 | Br_Fr_Cor_BA9 |
| Brain Hippocampus | Br_Hippoc |
| Brain Hypothalamus | Br_Hypothal |
| Brain Nucleus Accumbens Basal Ganglia | Br_Nucl_Accum_Bas_Gang |
| Brain Putamen Basal Ganglia | Br_Put_Bas_Gang |
| Brain Spinal Cord Cervical C-1 | Br_Spin_Cord_Cerv_C1 |
| Brain Substantia Nigra | Br_Subst_Nigra |
| Breast Mammary Tissue | Bre_Mam_Tis |
| Cells EBV-Transformed Lymphocytes | Cells_EBV-transf_lymph |
| Cells Transformed Fibroblasts | Cells_Transf_fibr |
| Colon Sigmoid | Colon_Sigm |
| Colon Transverse | Colon_Transv |
| Esophagus Gastroesophageal Junction | Esoph_Gastr_Junct |
| Esophagus Mucosa | Esoph_Mucosa |

| | |
|---|---|
| Esophagus Muscularis | Esoph_Musc |
| Heart Atrial Appendage | Heart_Atr_Append |
| Heart Left Ventricle | Heart_L_Ventr |
| Liver | Liver |
| Lung | Lung |
| Minor Salivary Gland | Minor_Saliv_Gland |
| Muscle Skeletal | Muscle_Skeletal |
| Nerve Tibial | Nerve_Tibial |
| Ovary | Ovary |
| Pancreas | Pancreas |
| Pituitary | Pituitary |
| Prostate | Prostate |
| Skin Not Sun Exposed Suprapubic | Skin_N_S_Exp_Supr |
| Skin Sun Exposed Lower Leg | Skin_S_Exp_Low_leg |
| Small Intestine Terminal Ileum | Small_Intes_Term_Ileum |
| Spleen | Spleen |
| Stomach | Stomach |
| Testis | Testis |
| Thyroid | Thyroid |
| Uterus | Uterus |
| Vagina | Vagina |
| Whole Blood | Whole_Blood |

### S7. Converge haplotypes

#### DNA sequencing

DNA was extracted from saliva samples using the Oragene protocol. A barcoded library was constructed for each sample. Sequencing reads obtained from Illumina Hiseq machines were aligned to Genome Reference Consortium Human Build 37 patch release 5 (GRCh37.p5) with Stampy (v1.0.17) [6] [5] [1] [1] [1] [2] [2] using default parameters, after filtering out reads containing adaptor sequencing or consisting of more than 50% poor quality (base quality <= 5) bases. Samtools (v0.1.18) [7] was used to index the alignments in BAM format [7] and Picardtools (v1.62) was used to mark PCR duplicates for downstream filtering. The Genome Analysis Toolkit's (GATK, version 2.6). Base quality score recalibration (BQSR) was then applied to the mapped sequencing reads using BaseRecalibrator in Genome Analysis Toolkit (GATK, basic version 2.6) [8] with the known insertion and deletion (INDEL) variations in 1000 Genomes Projects Phase 1 [9] and known single nucleotide polymorphisms (SNPs) from dbSNP (v137, excluding all sites added after v129) excluded from the empirical error rate calculation. GATKlite (v2.2.15) was then used to output sequencing reads with the recalibrated base quality scores while removing reads without the "proper pair" flag bit set by Stampy (1-5% of reads per sample) using the --read_filter ProperPair option (if the "proper pair" flag bit is set for a pair of reads, it means both reads in the mate-pair are correctly oriented, and their separation is within 5 standard deviations from the mean insert size between mate-pairs).

#### Variant calling, imputation, and phasing

Variant discovery and genotyping (for both SNPs and INDELs) at all polymorphic SNPs in 1000G Phase1 East Asian (ASN) reference panel[10] was performed simultaneously using post-BQSR sequencing reads from all samples using the GATK's UnifiedGenotyper (version 2.7-2-g6bda569). Variant quality score recalibration (VQSR) was then performed with GATK's VariantRecalibrator (v2.7-4-g6f46d11) in SNP variant calls using the SNPs in 1000 Genomes Phase 1 ASN Panel [9] as the known, truth and training sets. A sensitivity threshold of 90% to SNPs in the 1000G Phase1 ASN panel was applied for SNP selection for imputation after optimizing for Transition to Transversion (TiTv) ratios in SNPs called. Genotype likelihoods (GLs) were calculated at selected sites using a sample-specific binomial mixture model implemented in SNPtools (version 1.0), and imputation was performed at those sites without a reference panel using BEAGLE (version 3.3.2) [11]. The second round of imputation was performed with BEAGLE on the same GLs, but only at biallelic SNPs polymorphic in the 1000G Phase 1 ASN panel using the 1000G Phase 1 ASN haplotypes as a reference panel. The genotypes derived from Beagle imputation were phased using Shapeit (version 2, revision 790) [12]. Genetic maps were obtained from the Impute2 [13] website. Chromosomes 13 - 22 and X were phased using 12 threads and default parameters. Chromosomes 1-12 were phased using 12 threads in four chunks that overlap by 1MB. The phased chunks were ligated together using ligateHAPLOTYPES, available from the Shapeit website. A final set of allele dosages and genotype probabilities was generated from these two datasets by replacing the results in the former with those in the latter at all sites imputed in the latter. We then applied a conservative set of inclusion threshold for SNPs for genome-wide association study (GWAS): a) p-value for violation HWE > $10^{-6}$, b) Information score > 0.9, c) MAF in CONVERGE > 0.5% to arrive at the final set of 6,242,619 SNPs. Details can be found in [14].

### S8. Reference panel

The reference panel includes the publicly available 22,691 subjects from Haplotype Reference Consortium (HRC) and 10,262 CONVERGE. For CONVERGE subjects, we used the province of origin to divide them into 4 population (CNE, CCE, CSE and CCS). HRC subjects coming from the small Orkney (ORK) island provided the basis for an additional European population, i.e. ORK.1KG subject from HRC and subjects from CONVERGE and ORK along with their a) population label b) first 20 ancestry principal components were used to train a quadratic discriminant model. Subsequently, to have more homogeneous populations in the panel, all available subjects were reassigned to subpopulations by using model prediction (Table S3). Consequently, many subjects might have been re-assigned to a different (but related) population.

**Table S2. Subpopulations in the reference panel and their continental cohort (super population).** EUR is the abbreviation for Europeans, AFR for Africans, ASN for Asians, AMR for Americans and SAS for south Asians.

| Population Abbreviation | Number of Subjects | Super Population | Population Description |
|---|---|---|---|
| ACB | 164 | AFR | African Caribbeans in Barbados |
| ASW | 162 | AFR | African Ancestry in |

| | | | |
|---|---|---|---|
| | | | Southwest US |
| BEB | 86 | SAS | Bengali from Bangladesh |
| CCE | 3,409 | ASN | China Central East |
| CCS | 2,613 | ASN | China Central South |
| CDX | 95 | ASN | Chinese Dai in Xishuangbanna, China |
| CEU | 6,360 | EUR | Utah residents with Northern and Western European ancestry |
| CLM | 98 | AMR | Colombians from Medellin, Colombia |
| CNE | 2,330 | ASN | China North East |
| CSE | 2,020 | ASN | China South-East |
| ESN | 140 | AFR | Esan in Nigeria |
| FIN | 3,529 | EUR | Finnish in Finland |
| GBR | 2,020 | EUR | British in England and Scotland |
| GIH | 110 | SAS | Gujarati Indian from Houston, Texas |
| GWD | 113 | AFR | Gambian in Western Divisions of Gambia |
| IBS | 1,309 | EUR | Iberian Population from Spain |

| | | | |
|---|---|---|---|
| ITU | 95 | SAS | Indian Telugu from the UK |
| JPT | 107 | ASN | Japanese in Tokyo, Japan |
| KHV | 226 | ASN | Kinh in Ho Chi Minh City, Vietnam |
| LWK | 99 | AFR | Luhya in Webuye, Kenya |
| MSL | 87 | AFR | Mende in Sierra Leone |
| MXL | 187 | AMR | Mexican Ancestry from Los Angeles, USA |
| ORK | 5,772 | EUR | Orkney Island study |
| PEL | 110 | AMR | Peruvians from Lima, Peru |
| PJL | 121 | SAS | Punjabi from Lahore, Pakistan |
| PUR | 138 | AMR | Puerto Rican in Puerto Rico |
| STU | 110 | SAS | Sri Lankan Tamil from the UK |
| TSI | 1,291 | EUR | Tuscani in Italia |
| YRI | 52 | AFR | Yoruba in Ibadan, Nigeria |

### S9. Assessing the Type I error rate

To compare the Type I error rate of the proposed method JEPEGMIX2-P, we estimated the relative Type I error (the empirical divided by the nominal Type I error rate) as a function of the nominal Type I error rate, (shown on -log10 scale in Fig. S1-S5) for five cohorts. These cohorts were drawn according to five different cosmopolitan scenarios (Table S3) based on the 1000 Genomes haplotypic data: 1) 30% CEU + 25% CHS + 5% PUR + 40% YRI (Cohort 1), 2) 10% ASW + 15% CEU + 15% CHB + 12.5% CHS + 15% GBR + 10% MXL + 2.5% PUR + 20% YRI (Cohort 2), 3) 15% ASW + 35% CHB + 35% GBR + 15% MXL (Cohort 3), 4) 45% ASW + 55% GBR (Cohort 4) and 5) 55% CHB + 45% MXL (Cohort5). For each scenario, we obtained pathway statistics for two enrichment cases: i) under null ($H_0$) and ii) polygenic null ($H_p$) – where the entire genome is roughly equally enriched. Each scenario was analyzed with JEPEGMIX2-P assuming prespecified weights (PRE), i.e. weights were assigned based on study information, and automatically estimate weights (EST), i.e. those estimated by JEPEGMIX2-P.

To test the size of the test for JEPEGMIX2-P under “a randomly enriched scenario”, we create a null polygenic case ($H_p$) by adding two independent realizations of the null hypothesis. This is equivalent to an enriched GWAS having a) unit non-centrality of the chi-square distribution and b) enrichment that is independent of any possible functionality variants of genetic regions. We still call it a null scenario because it is not preferentially enriched in any functional category, including eQTLs and coding regions.

We noted from our preliminary results that MSigDB pathways with character length names less or equals to 8, as chr15q25, ch6p21 etc., include genes in high LD due to their clustering in the same chromosome band. For that reason, we also estimate the size of the test for all cohort scenarios, cases, and enrichment, as above, only for these high LD pathways. Additional to test how good the JEPEGMIX2-P adjust for pre-estimated weights from super populations, we estimated the size of the test for all the above scenarios (five different cohorts, 100 data sets each, $H_0$, $H_p$ cases and pathway with character length names less or equals to 8).

**Table S3. Weight declaration for Cohorts from 1000 Genomes haplotypic data.** CEU denotes Utah Residents (CEPH) with Northern and Western European Ancestry, CHB - Han Chinese in Beijing, China, CHS - Han Chinese in the South (China).

| Cohorts | ASW | CEU | CHB | CHS | GBR | MXL | PUR | YRI |
|---|---|---|---|---|---|---|---|---|
| Coh1 | | 0.30 | | 0.25 | | | 0.05 | 0.40 |
| Coh2 | 0.10 | 0.15 | 0.15 | 0.125 | 0.15 | 0.10 | 0.025 | 0.20 |
| Coh3 | 0.15 | | 0.35 | | 0.35 | 0.15 | | |
| Coh4 | 0.45 | | | | 0.55 | | | |
| Coh5 | | | 0.55 | | | 0.45 | | |

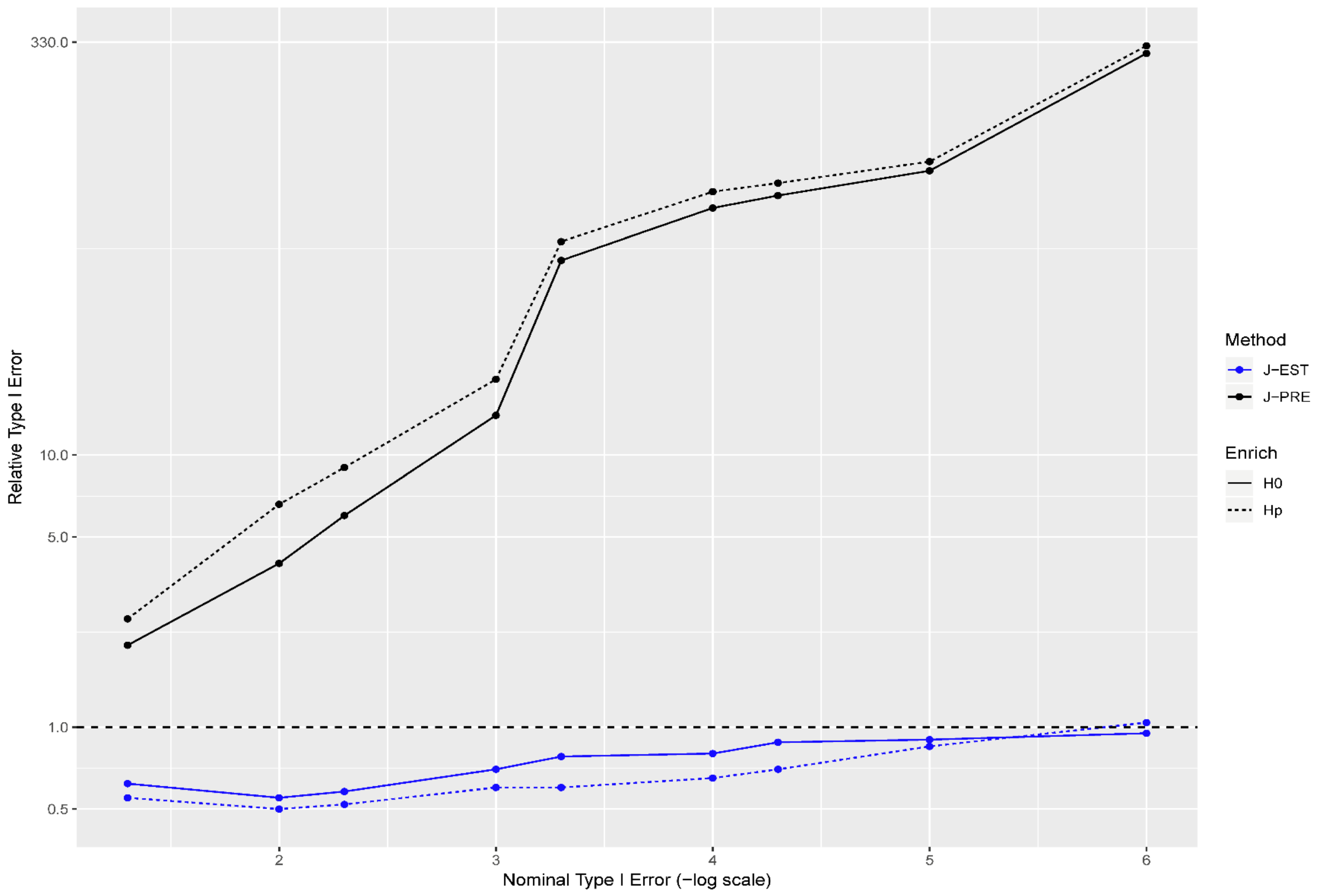


**Fig. S1. Relative size of the test (the quotient of empirical false positive rate and nominal type I error), for all pathways in the analysis of Cohort1 (30% CEU + 25% CHS + 5% PUR + 40% YRI). In legend, enrich designates whether the statistic was under null ($H_0$) or polygenic null ($H_p$) hypotheses. Method denotes whether the statistics had estimate weights (EST) or pre-estimate (PRE).**

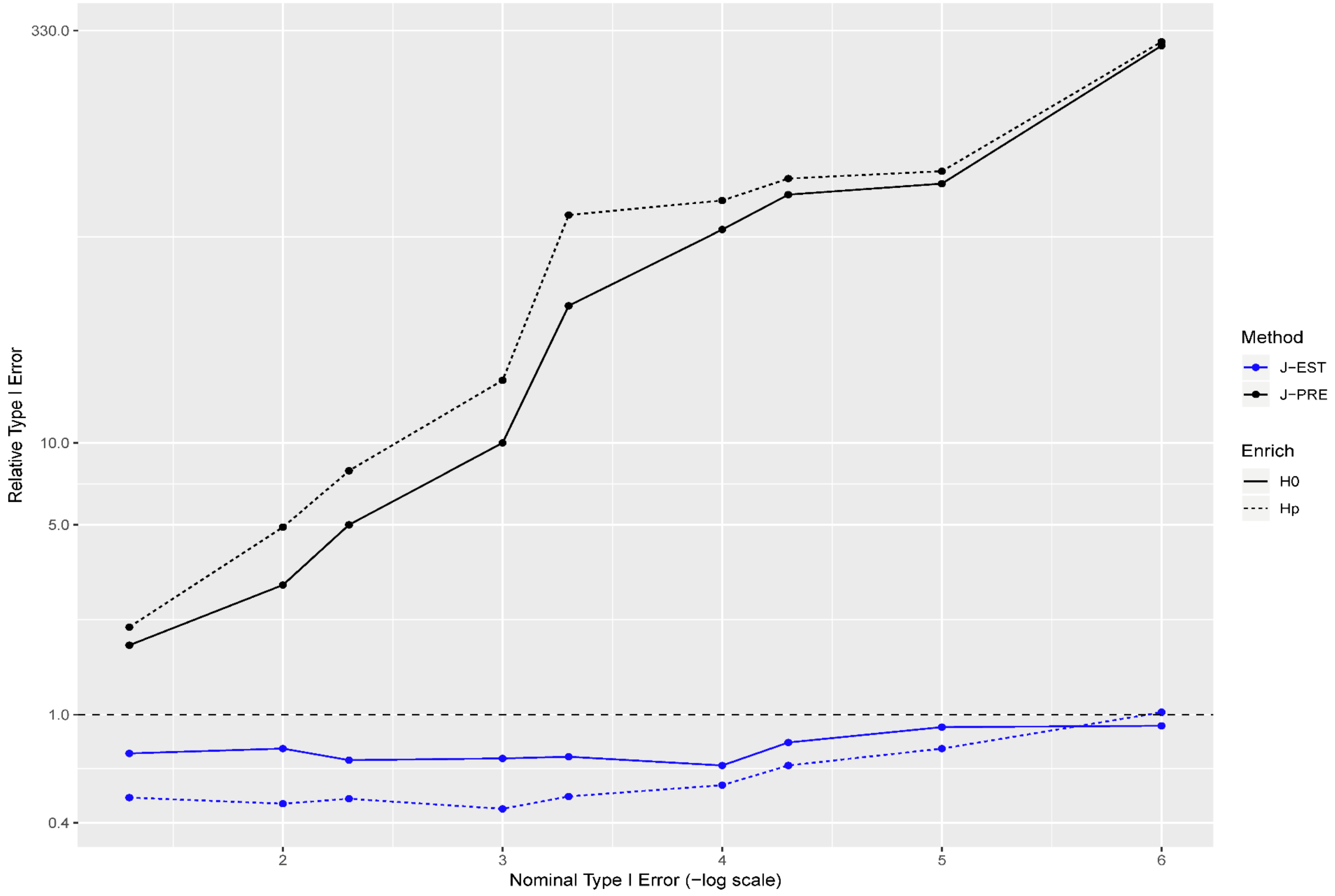


**Fig. S2. Relative size of the test for Cohort2 (10% ASW + 15% CEU + 15% CHB + 12.5% CHS + 15% GBR + 10% MXL + 2.5% PUR + 20% YRI). See Fig S1 for background and abbreviations.**

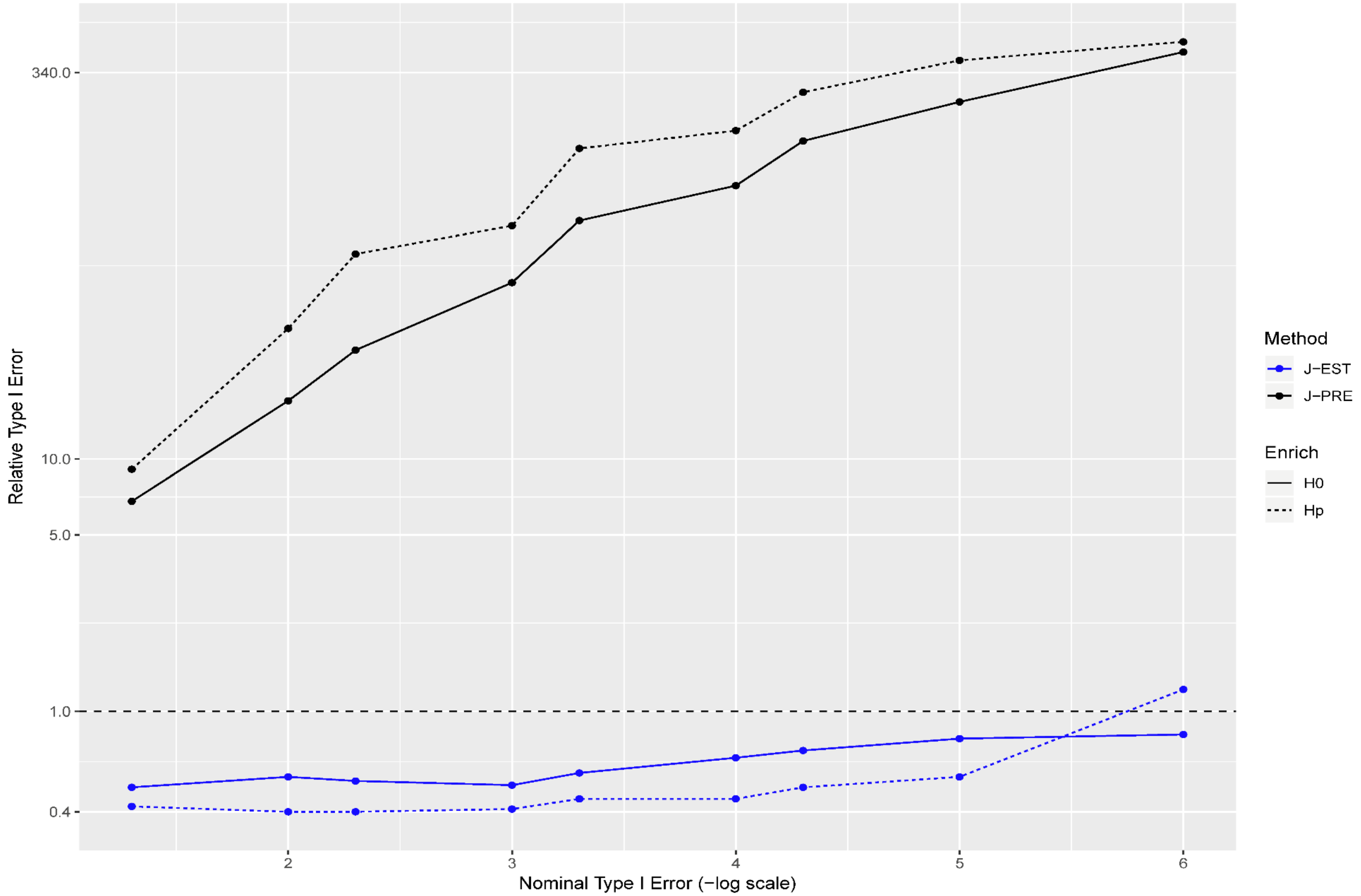


**Fig. S3. Relative size of the test for Cohort3 (15% ASW + 35% CHB + 35% GBR + 15% MXL). See Fig S1 for background and abbreviations.**

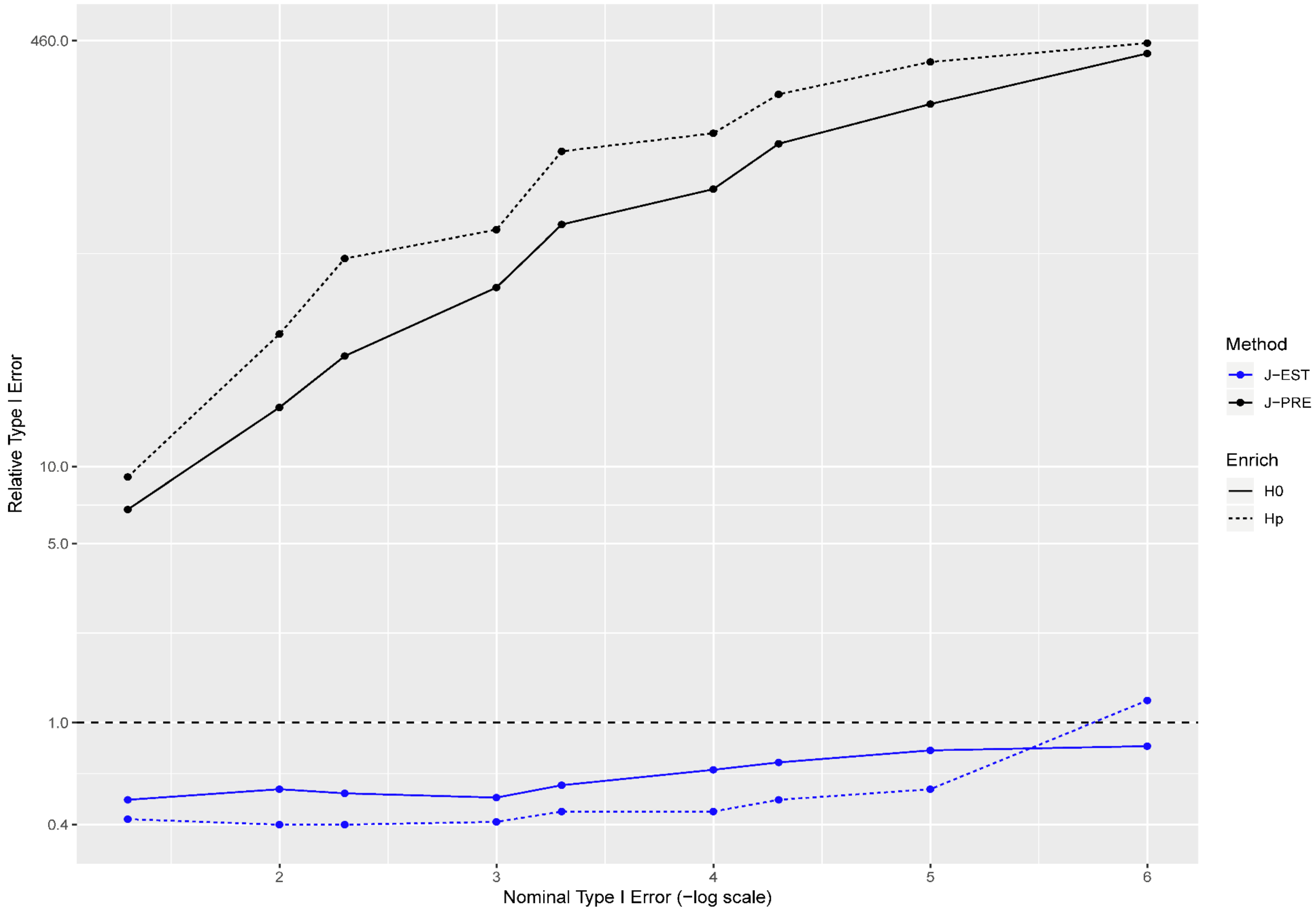


**Fig. S4. Relative size of the test for Cohort4 (45% ASW + 55% GBR). See Fig S1 for background and abbreviations.**

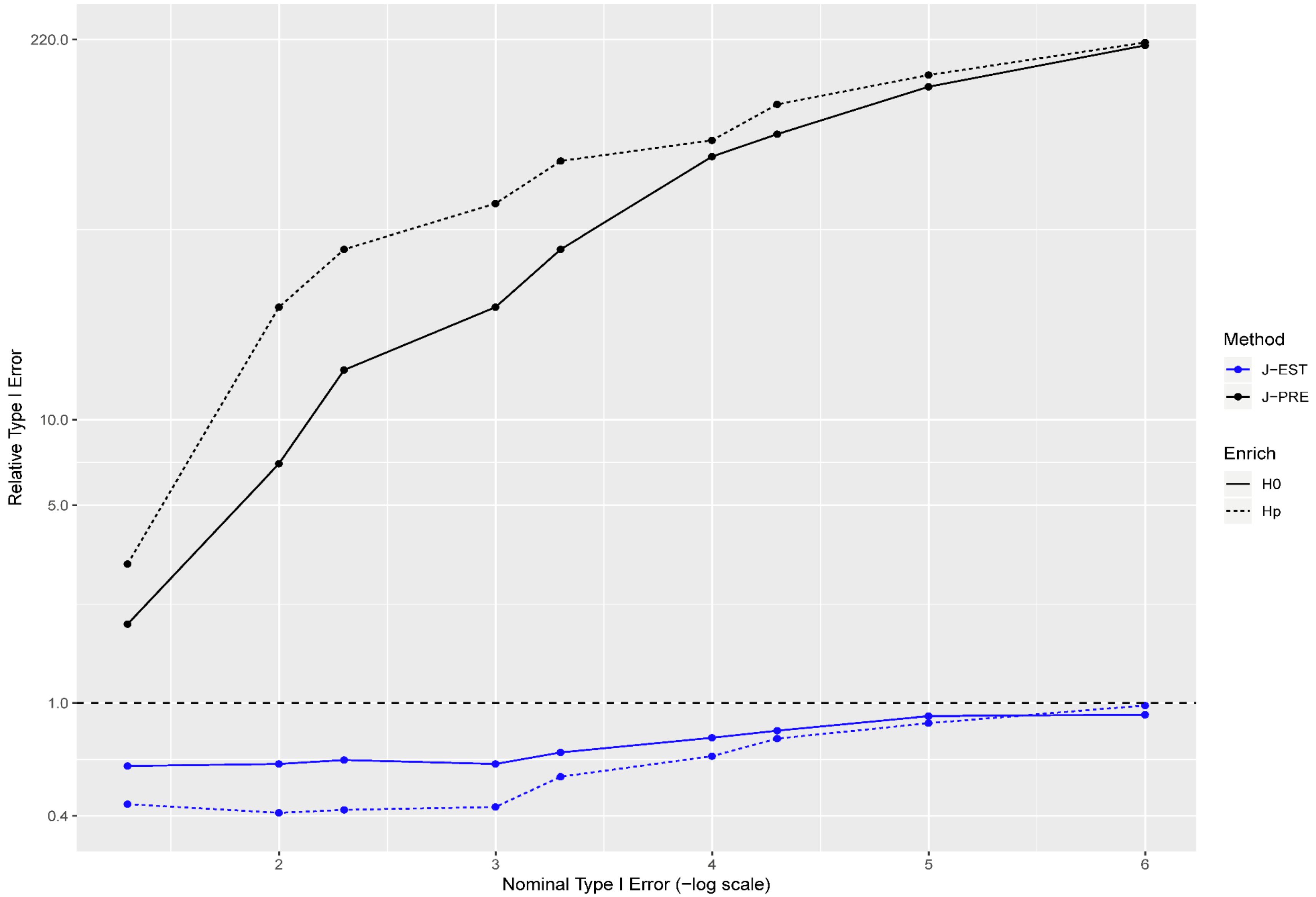


**Fig. S5. Relative size of the test for Cohort5 (55% CHB + 45% MXL). See Fig S1 for background and abbreviations.**

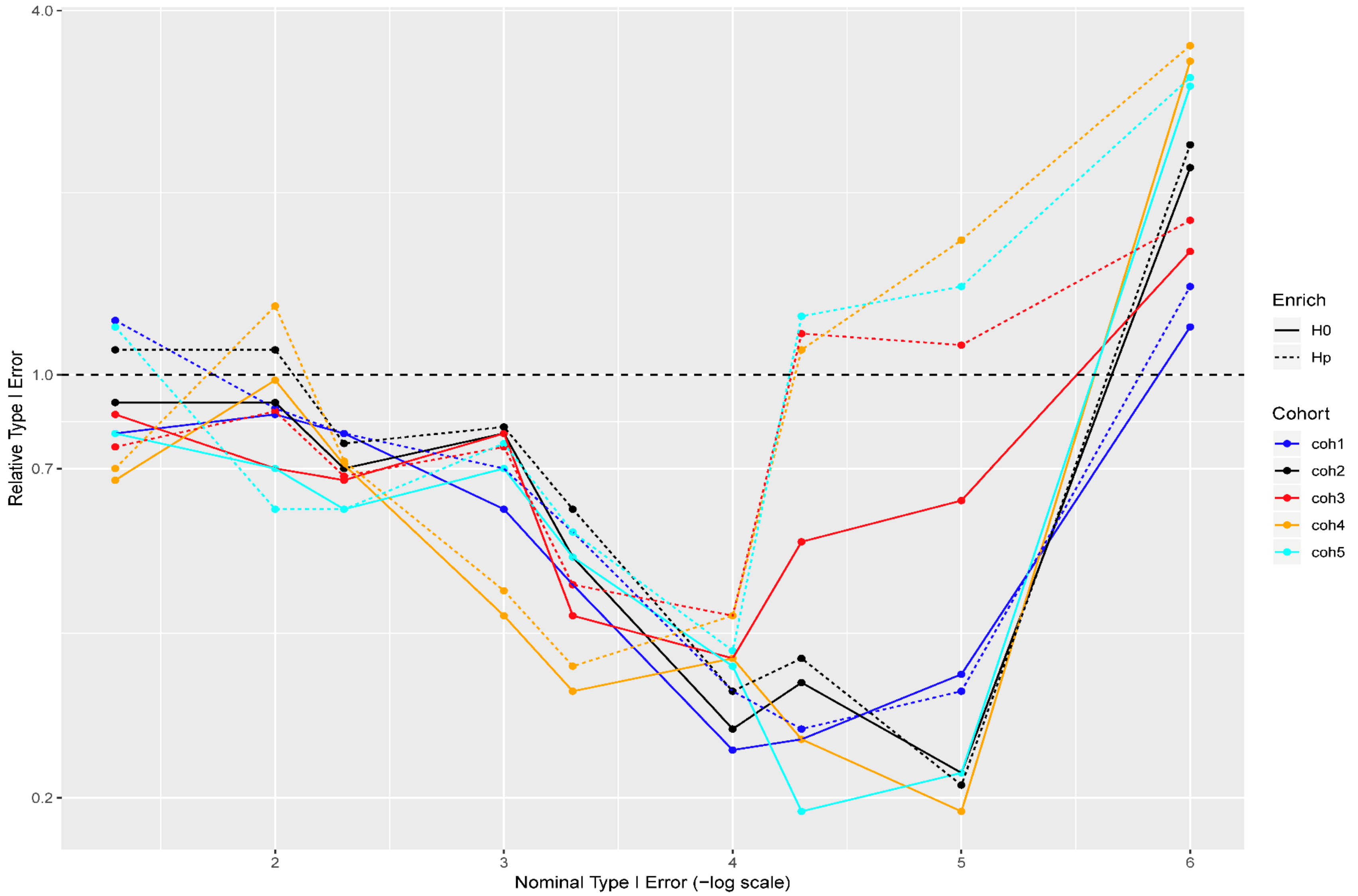


**Fig. S6. Relative size of the test, for all pathways in the analysis for all Cohort, when using pre-estimated weights according to the continental super populations (AFR, AMR, ASN, EUR, SAS). See Fig S1 abbreviations.**

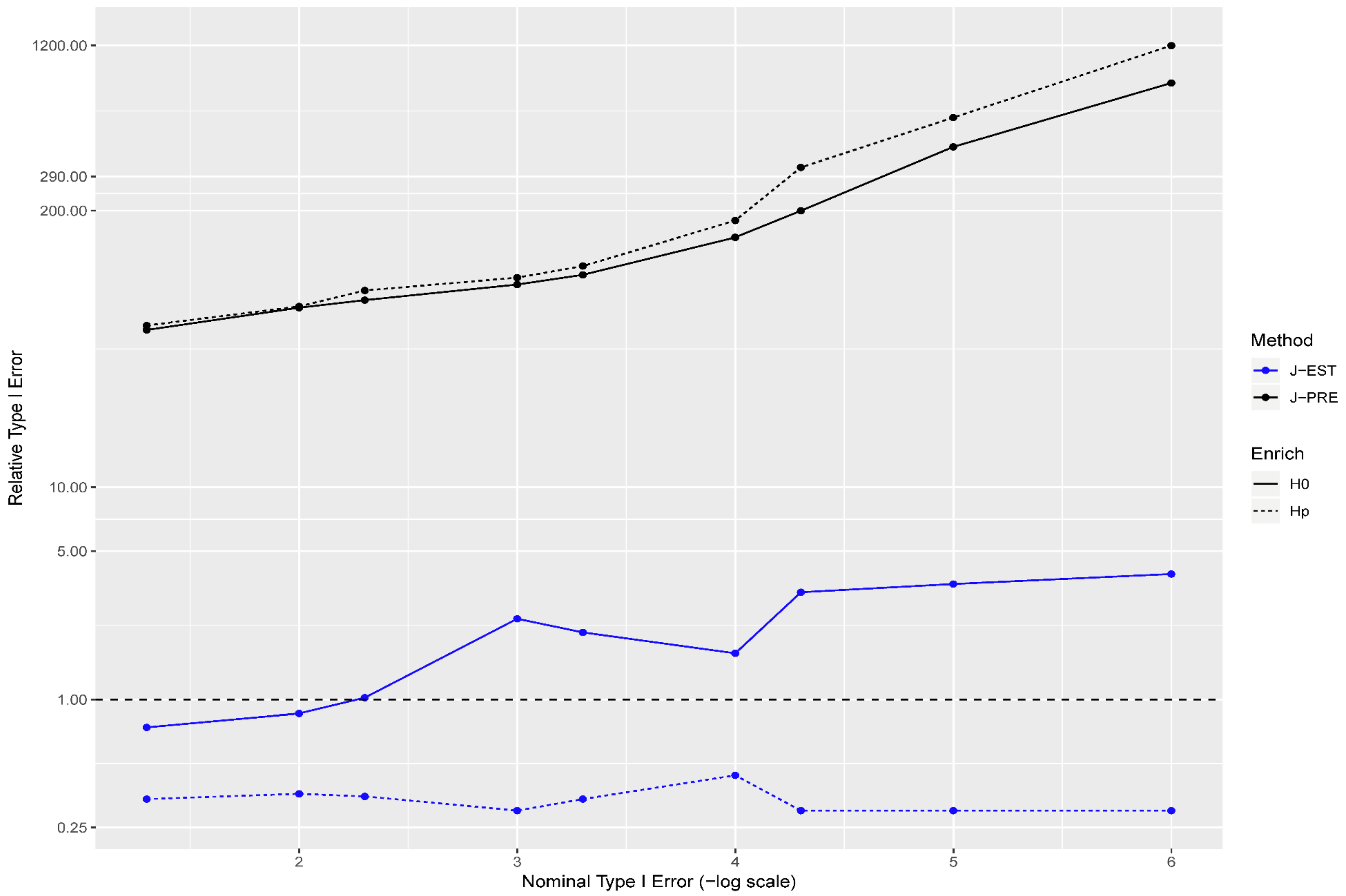


**Fig. S7. Relative size of the test for high-LD pathways (MSigDB pathways with name lengths $\leq 8$)- Cohort1 ancestry. See Fig S1 and S6 for background and abbreviations.**

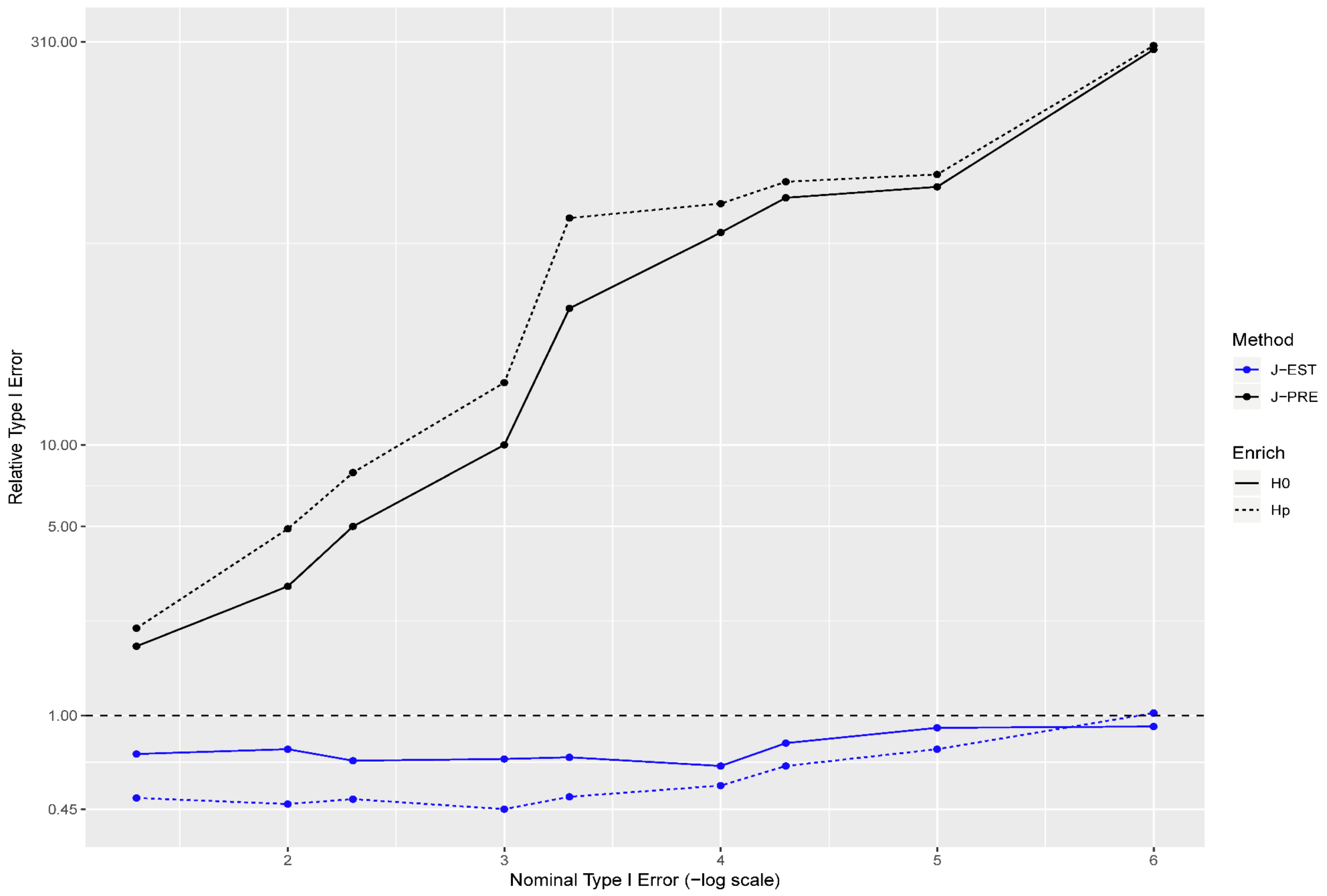


**Fig. S8. Relative size of the test for high-LD pathways - Cohort2. See Fig S1 and S7 for background and abbreviations.**

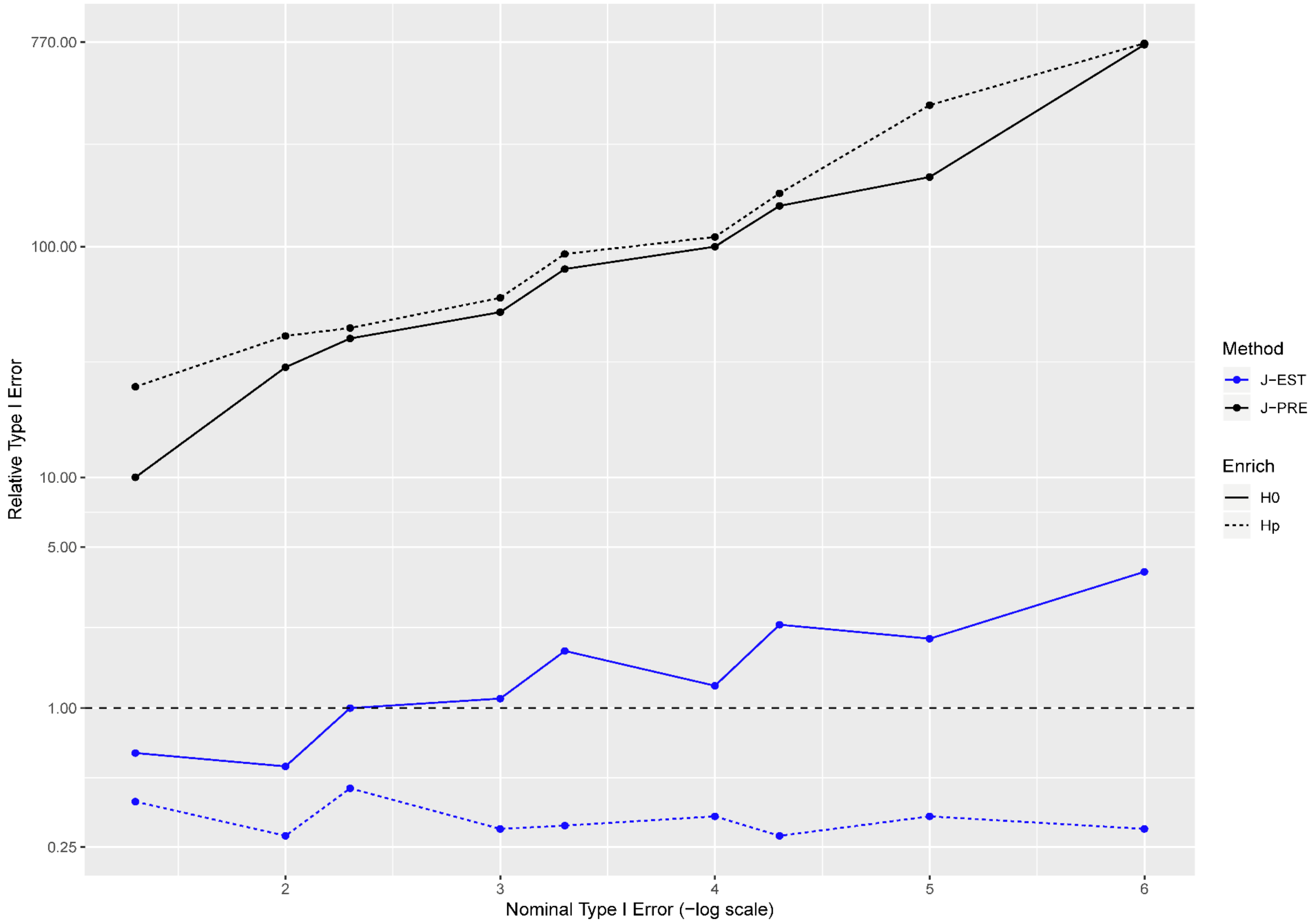


**Fig. S9. Relative size of the test for high-LD pathways - Cohort3. See Fig S1 and S7 for background and abbreviations.**

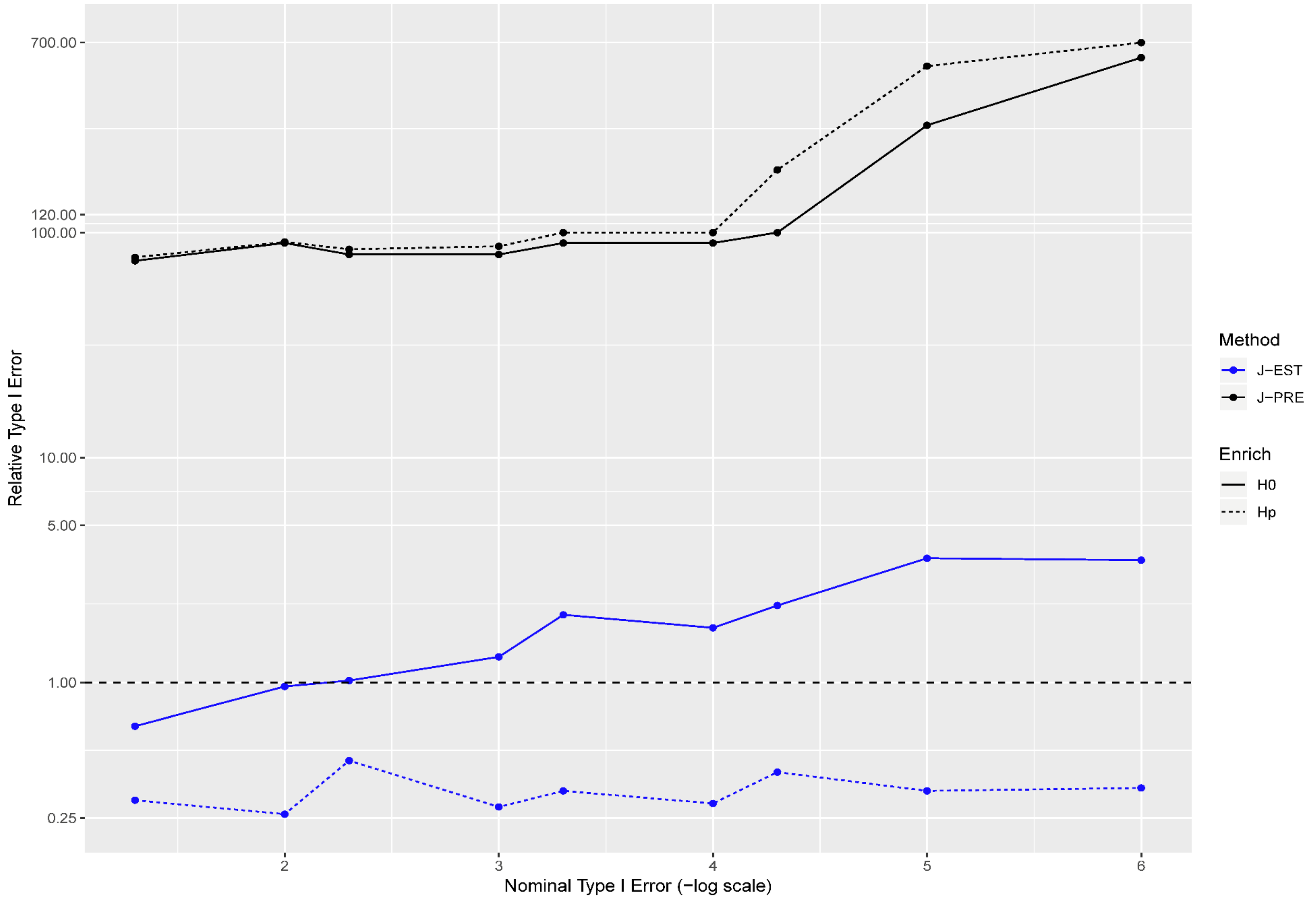


**Fig. S10. Relative size of the test for high-LD pathways - Cohort4. See Fig S1 and S7 for background and abbreviations.**

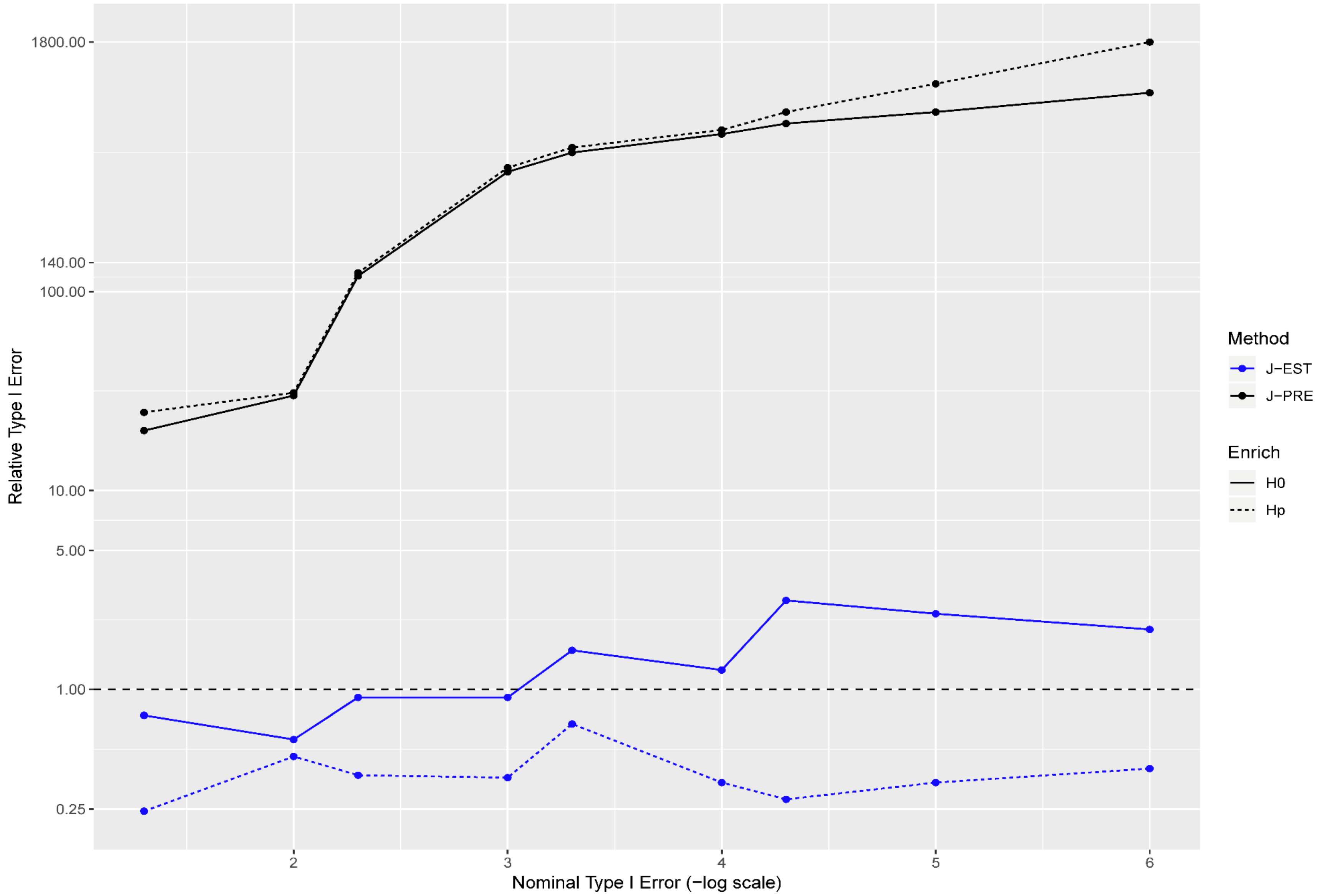


**Fig. S11. Relative size of the test for high-LD pathways - Cohort5. See Fig S1 and S7 for background and abbreviations.**

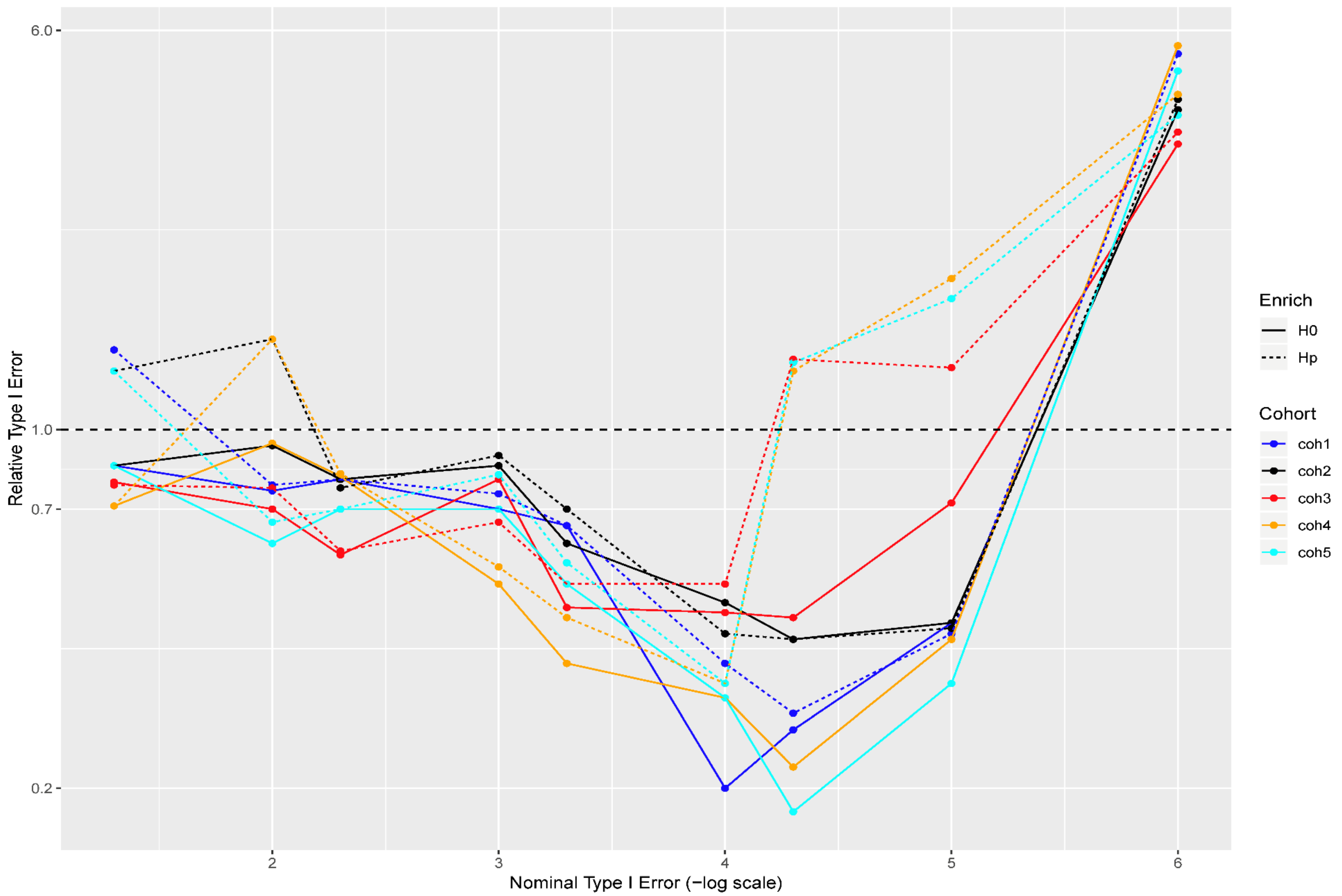


**Fig. S12 Relative size of the test for high-LD pathways in nullified data sets. See Fig S1, S6 and S7 for background and abbreviations.**

### S10. Practical Applications.

In this section, we obtained pathway-level statistics by applying JEPEGMIX2-P to association summary statistics from PGC: i.e. ADHD, AUT, BIP, ED, MDD2 and SCZ. For the above datasets, for JEPEGMIX2-P, we construct heatmaps for the significant pathways ($q < 0.05$) (Fig. S13-S19).

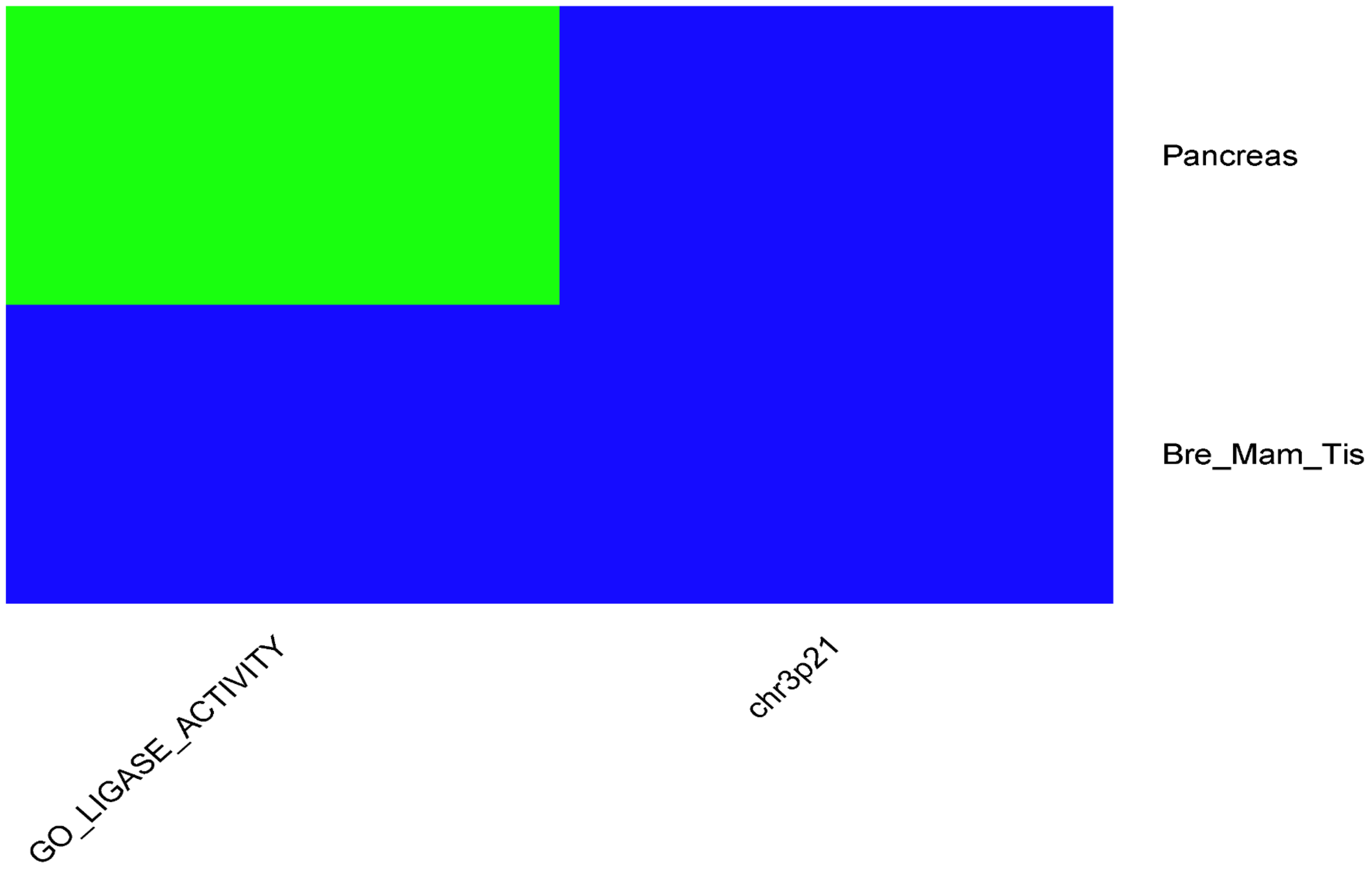


**Fig. S13.** ADHD pathway signals heatmap (conditional analysis yields similar map). The pathways (x-axis) and tissues (right hand y-axis) are ordered in the decreasing order of the overall sum of $-log10(p$-values) for all tissues and pathways with at least two significant signals. Where red color denotes $q < 0.001$, orange $0.001 < q < 0.01$, green $0.01 < q < 0.05$, light blue $0.05 < q < 0.16$ and blue $0.16 < q < 1$.

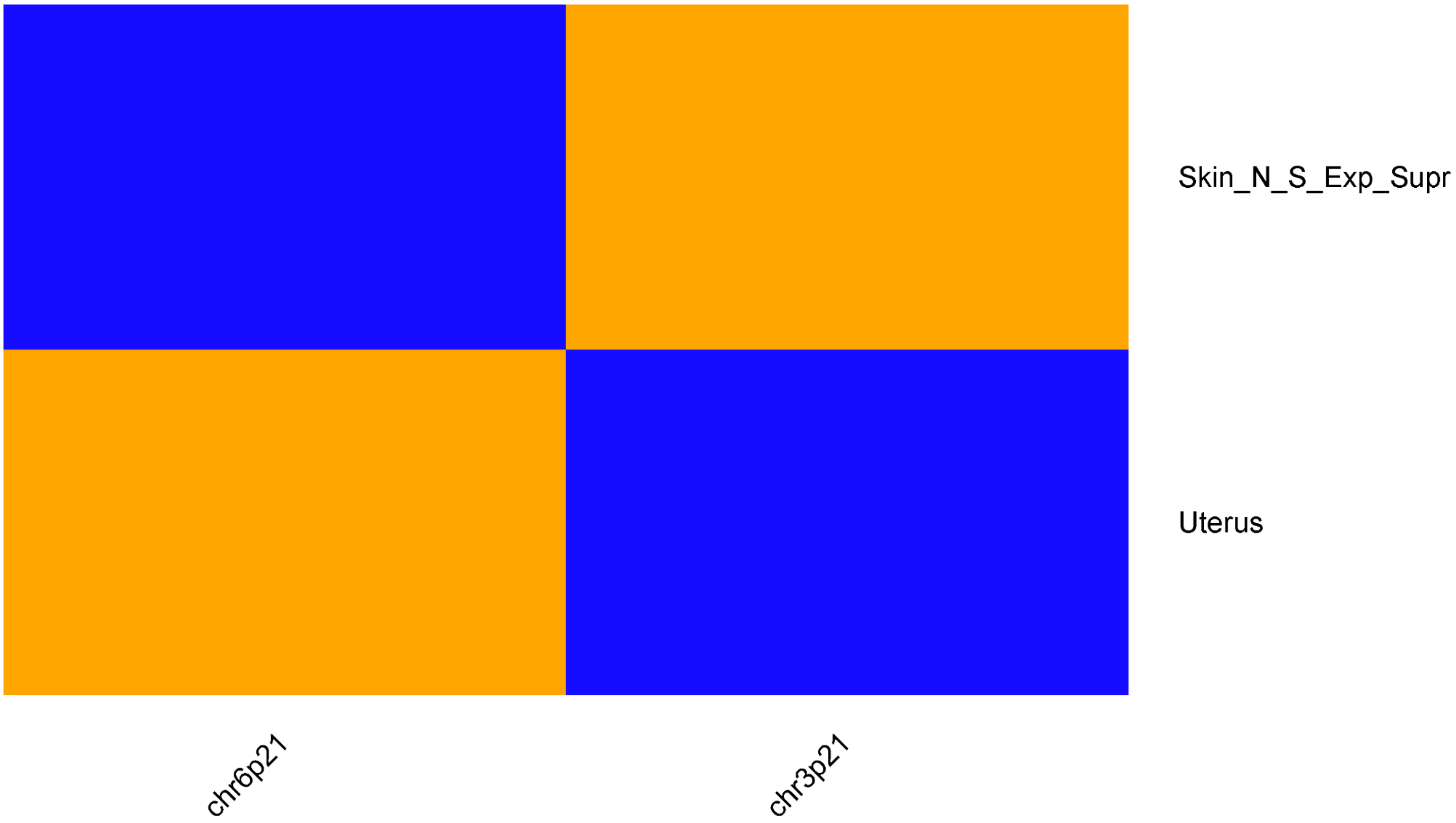


**Fig. S14 AUT pathway signals (conditional analysis yields no signal) heatmap. See Fig S13 for background.**

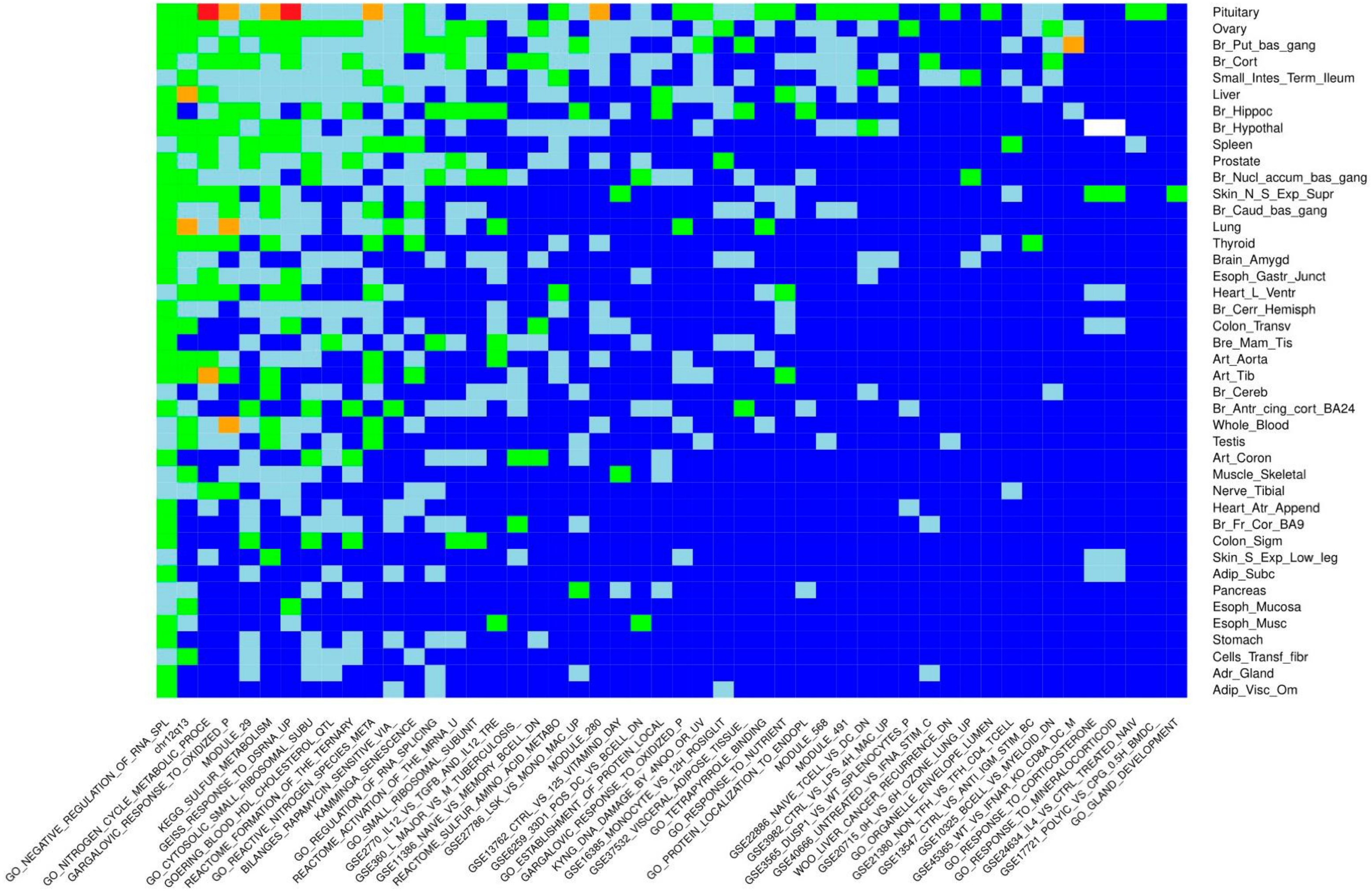


**Fig. S15 Top 50 ED (Anorexia) pathway signals heatmap (conditional analysis yields similar map). See Fig S13 for background.**

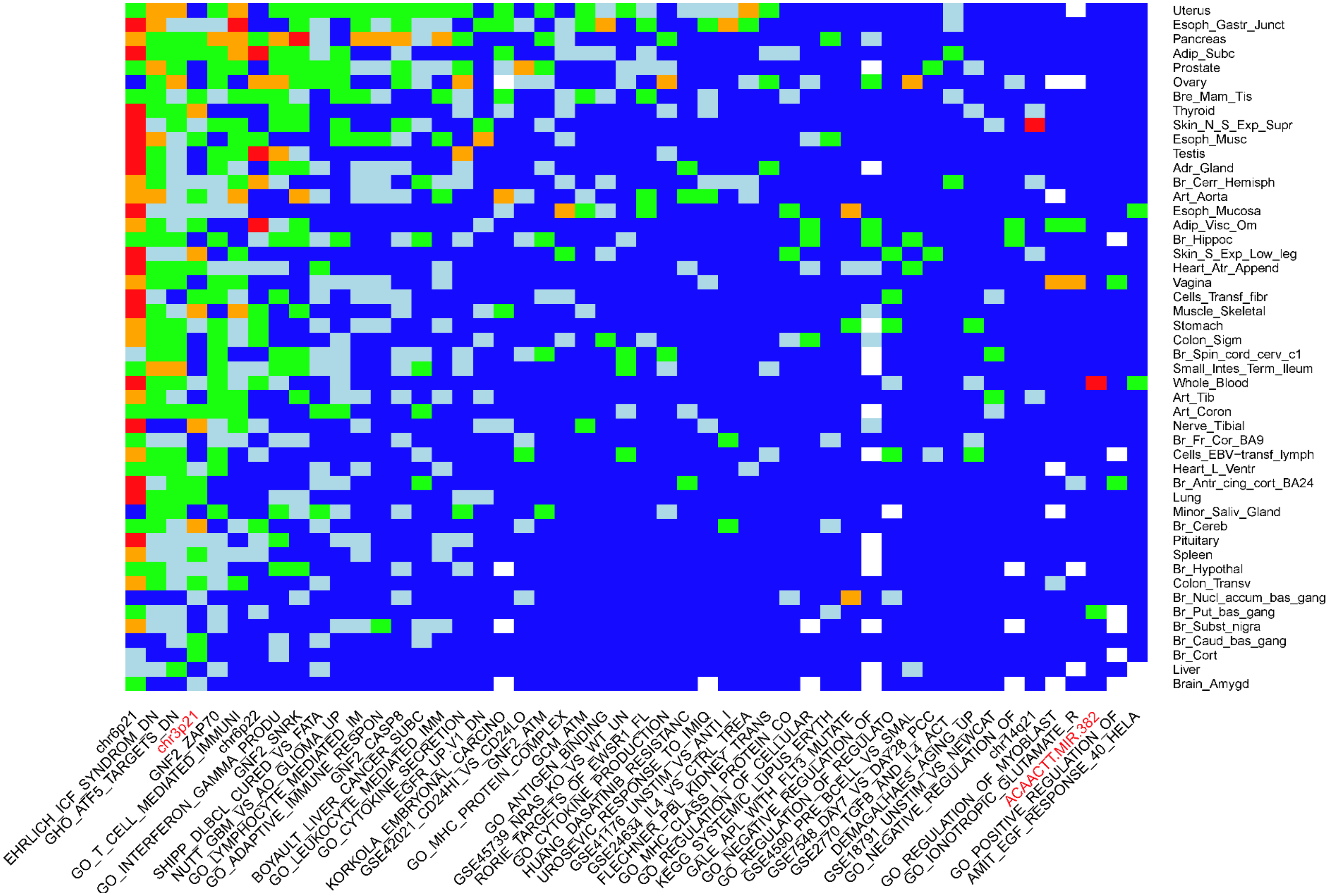


**Fig. S16 Top 50 MDD pathway unconditional signals heatmap. See Fig S13 for background.**

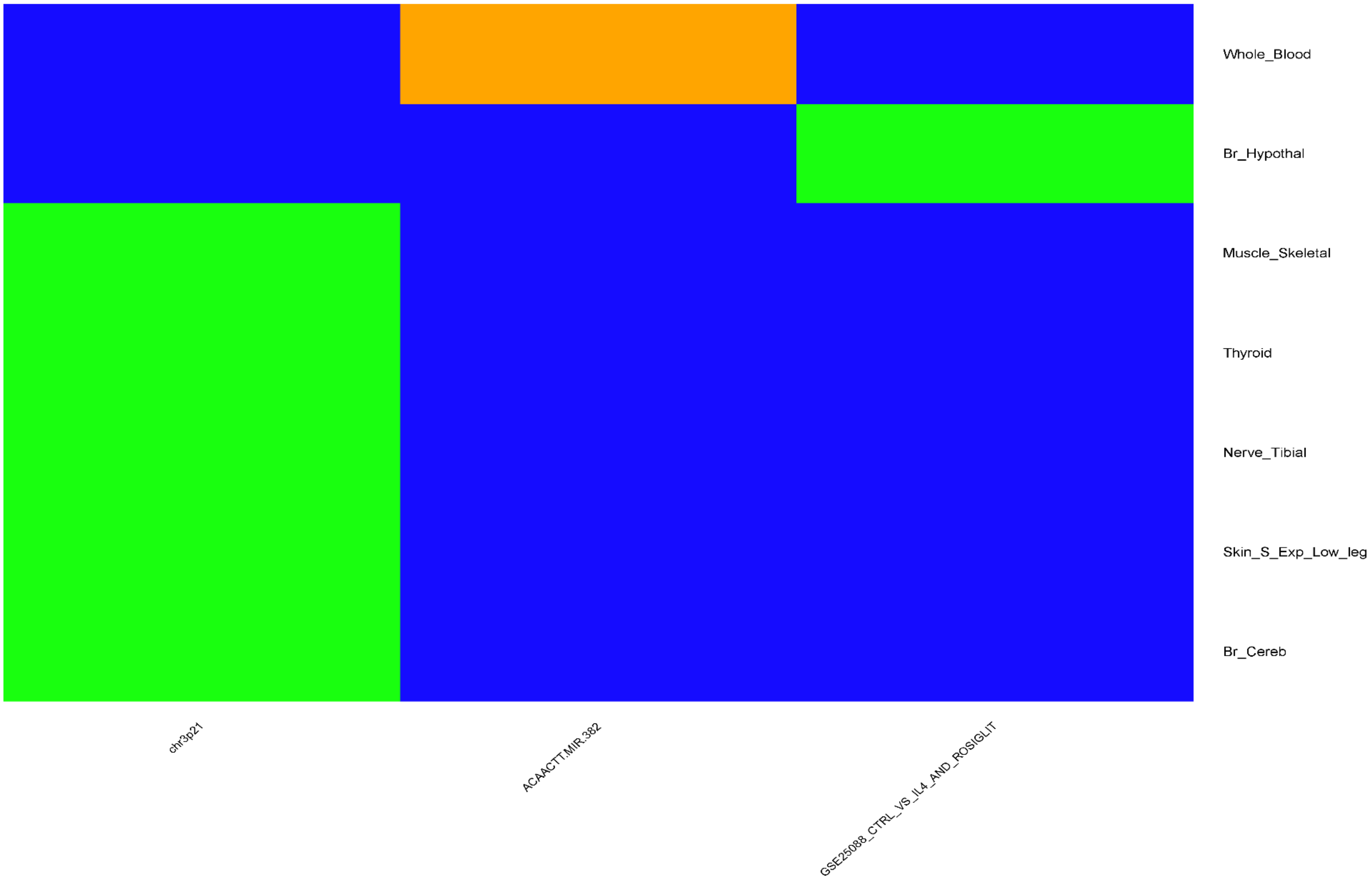


**Fig. S17 MDD heatmap for pathway signals after conditioning on significant SNP signals. See Fig S13 for background.**

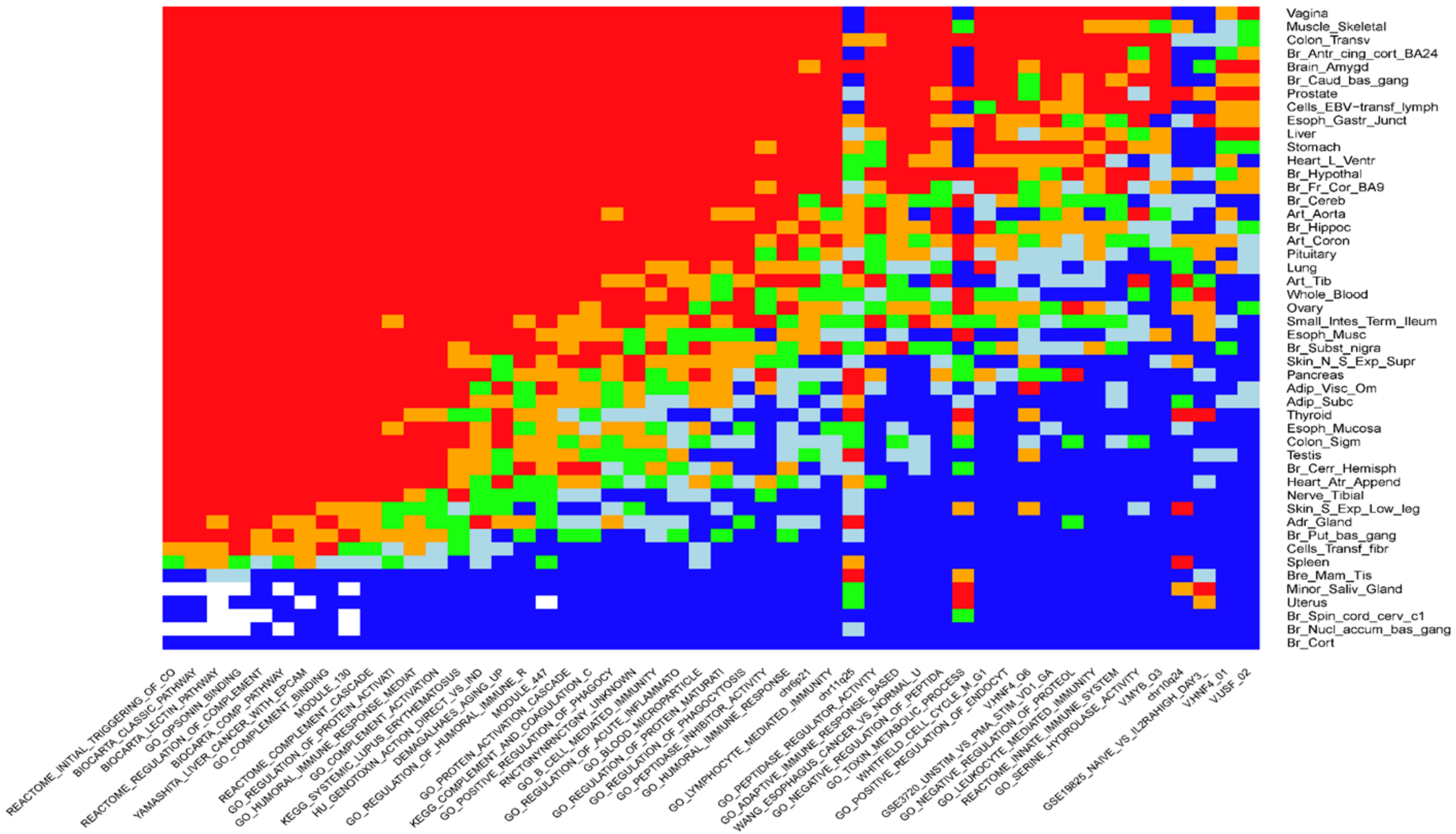


**Fig. S18 Top 50 SCZ pathway unconditional signals heatmap. See Fig S13 for background.**

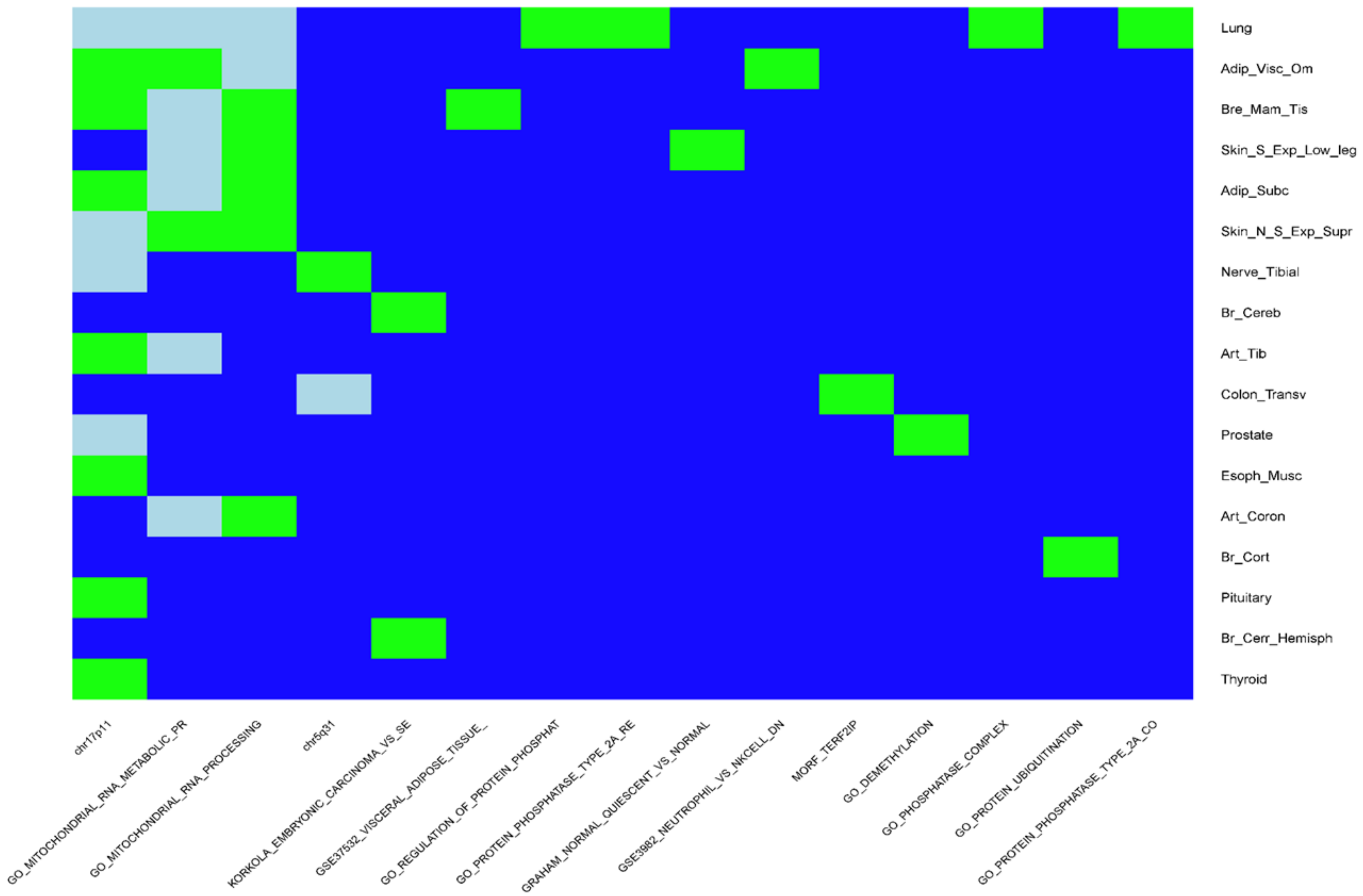


**Fig. S19 SCZ heatmap for pathway signals after conditioning on significant SNP signals. See Fig S13 for background.**
